# The ω subunit stabilizes transcribing RNA polymerase to balance processivity and collision resolution

**DOI:** 10.64898/2026.05.17.725479

**Authors:** Barbare Khitiri, Bing Wang, Matthew B. Cooke, Luca Buccolieri, David Dulin, Christophe Herman, Irina Artsimovitch

## Abstract

The ubiquitous subunit of RNA polymerase (RNAP), ω/RPB6, is traditionally viewed as an assembly chaperone or bacterial σ-factor competition modulator. This study redefines the role of *Escherichia coli* ω, encoded by the *rpoZ* gene. Unexpectedly, *ΔrpoZ* strain does not exhibit major defects in σ^S^-dependent stress responses, indicating its primary function lies elsewhere. Our CRISPRi screen suggested that losing ω may promote survival during transcription-replication conflicts. Consistently, we show that loss of ω sensitizes RNAP to termination, reduces RNAP processivity, and suppresses toxic effects of DNA-damaging agents in strains lacking functional DksA, Rho, or SeqA; DksA and Rho promote the release of stalled RNAP from nucleic acids, while SeqA prevents aberrant replication initiation. These findings suggest that loss of ω facilitates the removal of stalled RNAP, preventing catastrophic replisome collisions. We propose that ω/RPB6 homologs may balance RNAP processivity with controlled release to preserve genome integrity across all domains of life.

## Introduction

In all cells, transcription is carried out by multi-subunit RNA polymerase (RNAP) enzymes that share an ancestral origin, a crab-claw architecture, and structural domains that mediate RNAP interactions with nucleic acids and catalysis (*1*). Although RNAP composition varies across life, a “minimal” conserved core enzyme consists of five subunits, α^Ι^, α^II^, β, β’, and ω in bacteria (*1, 2*). The α^Ι^, α^II^, β, and β’ subunits are essential for catalysis and regulation, but the role of the smallest ω subunit remains elusive (*3*).

Availability of RNAP structures led to a seminal finding that ω and yeast RPB6 “latch” onto the β’/RPB1 subunit, assisting RNAP assembly and perhaps stabilizing its structure (*4*). Curiously, while its eukaryotic and archaeal counterparts are essential, ω is dispensable in bacteria, possibly because a general chaperone GroEL can promote RNAP assembly in the absence of ω (*3*).

The ω subunit is encoded in nearly every sequenced bacterial genome (**Fig. 1A, Table S1**) and interacts with the C-terminus of the catalytic β’ subunit in transcription complex structures (**Fig. 1B**). The three helices of the N-terminal of ω homologs from distant phyla fold similarly (**Fig. S1**). However, the genomic context of the gene that encodes ω (*rpoZ* in *Escherichia coli*; **Fig. 1C**) and the extent of β’/ω contacts (**Fig. 1B**) vary significantly in different bacterial phyla.

**Fig. 1.**
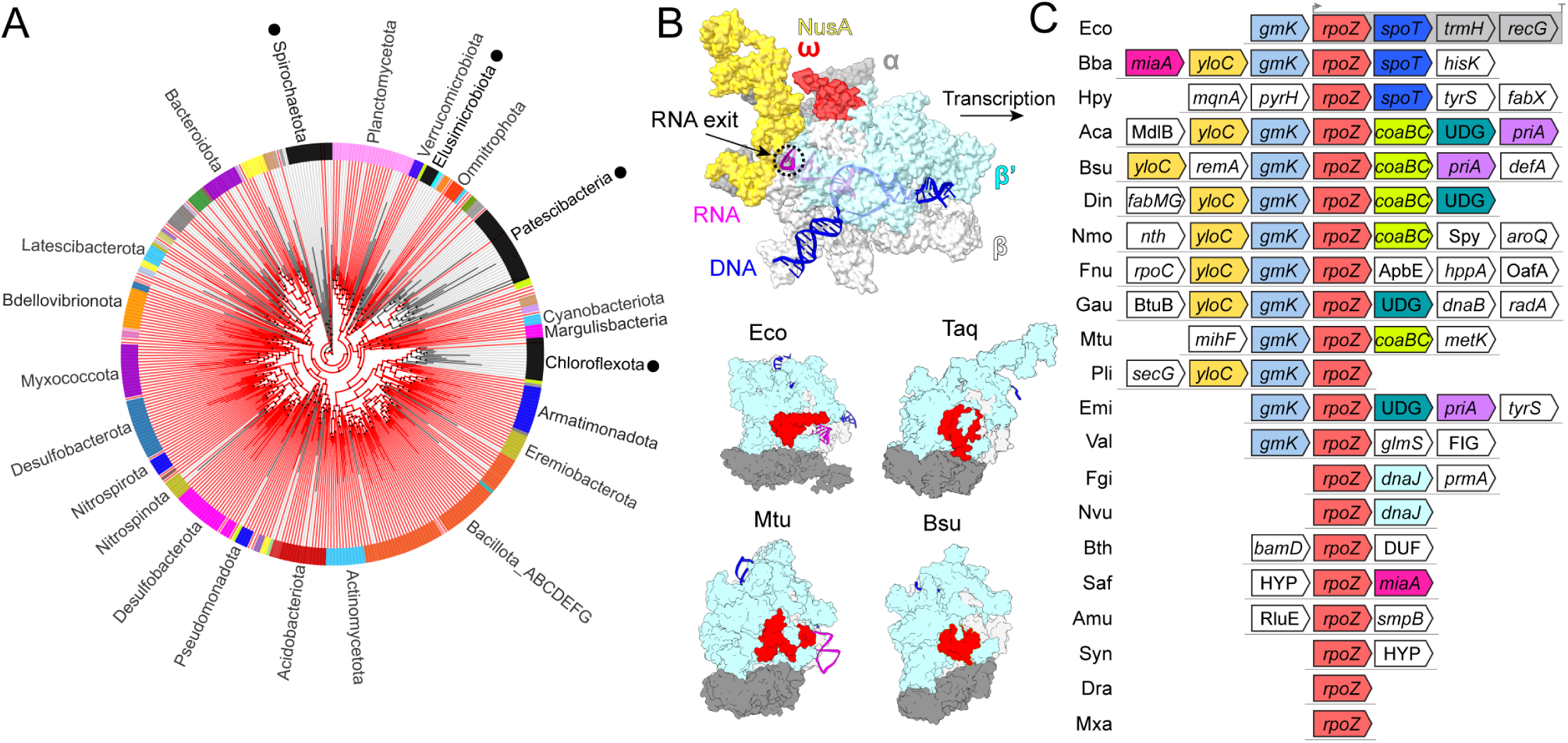
The distribution of RNAP subunit ω homologs in the bacterial kingdom. (**A**) The RNAP ω subunit is present in most genomes; the phylogenetic tree is viewed at the class level. The absence of ω may be a false negative: ω is small, poorly conserved, and is encoded in different neighborhoods; we thus manually examined the four phyla apparently lacking ω (indicated by black dots; **Table S1**). (**B**) The structure of *E. coli* RNAP transcription elongation complex (PDB ID: 6FLQ) is shown as a transparent surface. Four structures (only core subunits are shown) from Eco (*E. coli*; PDB ID: 6FLQ), Taq (*Thermus aquaticus*; PDB ID: 5TJG), Mtu (*Mycobacterium tuberculosis*; PDB ID: 8E79), and Bsu (*Bacillus subtilis*; PDB ID: 6WVJ) showing the conserved binding site of ω. The figure was prepared with ChimeraX-1.9. (**C**) Co-directional gene neighbors of *rpoZ* homologs. Representatives from 21 phyla were investigated. A 200 bp cut-off was used to define neighboring genes. If a long operon is adjacent to *rpoZ*, only the three closest genes are shown. If a gene name/function is unknown, the identified domain is listed without italic font. Genes are not on scale. UDG, uracil-DNA glycosylase. DUF, Domain of Unknown Function. HYP, hypothetical protein. *hisK*, histidine kinase. Bba, *Bdellovibrio bacteriovorus*; Hpy, *Helicobacter pylori;* Aca, *Acidobacterium capsulatum;* Din, *Desulfurispirillum indicum*; Nmo, *Nitrospira moscoviensis*; Fnu, *Fusobacterium nucleatum;* Gau, *Gemmatimonas aurantiaca*; Pli, *Planctopirus limnophila*; Emi, *Elusimicrobium minutum*; Val, *Vulcanimicrobium alpinum*; Fgi, *Fimbriimonas ginsengisoli Gsoil*; Nvu, *Nitratidesulfovibrio vulgaris*; Bth, *Bacteroides thetaiotaomicron*; Saf, *Spirochaeta africana*; Amu, *Akkermansia muciniphila*; Syn, *Synechocystis* sp. PCC 6803; Dra, *Deinococcus radiodurans*; Mxa, *Myxococcus xanthus*.

Although bacterial strains lacking ω display only mild growth defects, its ubiquity sustained a search for additional regulatory roles of ω (*3*). Studies in *E. coli*, *Staphylococcus aureus*, and *Synechocystis* sp. suggested that ω modulates promoter selection by favoring RNAP interactions with the primary σ factor to promote the expression of housekeeping genes (*5–7*). This model is supported by gene expression profiling and analyses of RNAP composition: the loss of ω led to increased expression of genes controlled by alternative σ factors, such as the general stress response σ^S^ in *E. coli*, and to depletion of the primary σ from the cellular holoenzyme pools (*5–7*). These findings are consistent with the ω proximity to the β flap domain, which interacts with σ in promoter complexes (*8, 9*), but cannot explain why ω homologs are essential for transcription by archaeal and eukaryotic RNAPs.

Two studies in *E. coli* suggested that ω may shape global gene expression beyond promoter selection.

First, in the Δ*rpoZ* strain, the increase in σ^S^-dependent transcription was linked to the genomic DNA relaxation, an inference supported by changes in plasmid DNA topology (*7*). Second, the possibility that ω may be allosterically connected to the RNAP active site was suggested by the identification of a dominant-lethal *rpoZ* N60D allele that was suppressed by the substitution at β’ Y457 (*10*), which is adjacent to the catalytic triad [β’ D460, D462, D464].

These reports, together with our observations that ω may be locked in an inactive state by a non-native C-terminal extension of the β’ subunit (*11*), prompted us to re-examine the ω contribution to cellular physiology. Surprisingly, despite the widely accepted role of ω in σ competition, we did not observe expected phenotypes in strains lacking ω. Here, we show that the precise deletion of the *rpoZ* gene in the *E. coli* MG1655 strain does not lead to the activation of the σ^S^ regulon or alter plasmid DNA topology. Instead, our results demonstrate that ω-less RNAP displays defects in processivity *in vivo* and *in vitro* and promotes *E. coli* survival under conditions that exacerbate the deleterious outcomes of transcription-replication collisions (TRCs). We show that the *rpoZ* deletion suppresses the toxic effects of DNA-damaging agents in strains without functional Rho or DksA, transcription factors that promote RNAP release from nucleic acids, or lacking SeqA, a DnaA inhibitor which prevents runaway DNA replication. Our findings support the original model in which ω functions as a ‘latch’ to stabilize the RNAP during and perhaps after its assembly (*4*), but redefine ω as a double-edged sword. The same’latch’ that ensures RNAP integrity becomes a molecular shackle when the polymerase stalls, turning into an immovable roadblock. Crucially, however, this roadblock is not absolute. The inherent ability of ω to fall off the stable core α2ββ’ assembly may provide a programmed release mechanism, allowing RNAP to shed its stabilizer and dissociate from the DNA template, letting the replisome go through.

## Results

### The rpoZ deletion does not activate the rpoS regulon

Previous studies were carried out with the *E. coli rpoZ* gene disrupted by a kanamycin-resistant cassette, which could have polar effects on the downstream genes that encode (p)ppGpp synthesize/dehydratase SpoT and helicase RecG (**Fig. 1C**). Indeed, the slow-growth phenotype of the *rpoZ::Kan^R^* strain was attributed to changes in the expression of SpoT, which regulates the amounts of the stringent response alarmone (p)ppGpp (*12*). In turn, changes in (p)ppGpp levels are known to modulate σ competition (*13*) and the chromosomal DNA supercoiling (*14*). RecG unwinds a broad range of substrates, co-localizes with the replisome (*15*), and has been implicated in plasmid replication control (*16*).

To avoid potential polar effects, we used the no-SCAR protocol, which combines λ-Red recombineering with CRISPR-Cas9 selection (*17*), to construct a “clean” *rpoZ* deletion in *E. coli* MG1655. We then performed a RNA-seq survey to compare the transcriptomes of the WT and *ΔrpoZ* strains, using a single replicate of each (**Fig. 2A**). This analysis identified 41 upregulated and 67 downregulated genes in the Δ*rpoZ* mutant (log₂FC > 2 or <-2, respectively). Based on studies in *E. coli* and other bacteria (*5–7*), we expected many upregulated genes to belong to the extensive RpoS regulon, which comprises nearly a quarter of *E. coli* genes. However, using a published MG1655 Δ*rpoS* RNA-seq dataset (*18*), we found no correlation in the gene expression between the Δ*rpoZ* and Δ*rpoS* strains (Pearson correlation of 0.13; **Fig. 2B**). To validate this finding, we used RT-qPCR to test the differential expression of three RpoS-dependent genes: *dps*, *katE*, and *osmE* (*19*). Consistent with our RNA-seq data, *rpoZ* deletion does not affect the expression of these genes (**Fig. 2C**). Additionally, we don’t observe ω-dependent changes in the topology of two different plasmids (**Fig. S2**), including pACYC184 that was used in a previous study (11). We suspect this discrepancy stems from polar effects caused by the *rpoZ* disruption on downstream genes in earlier work.

**Fig. 2.**
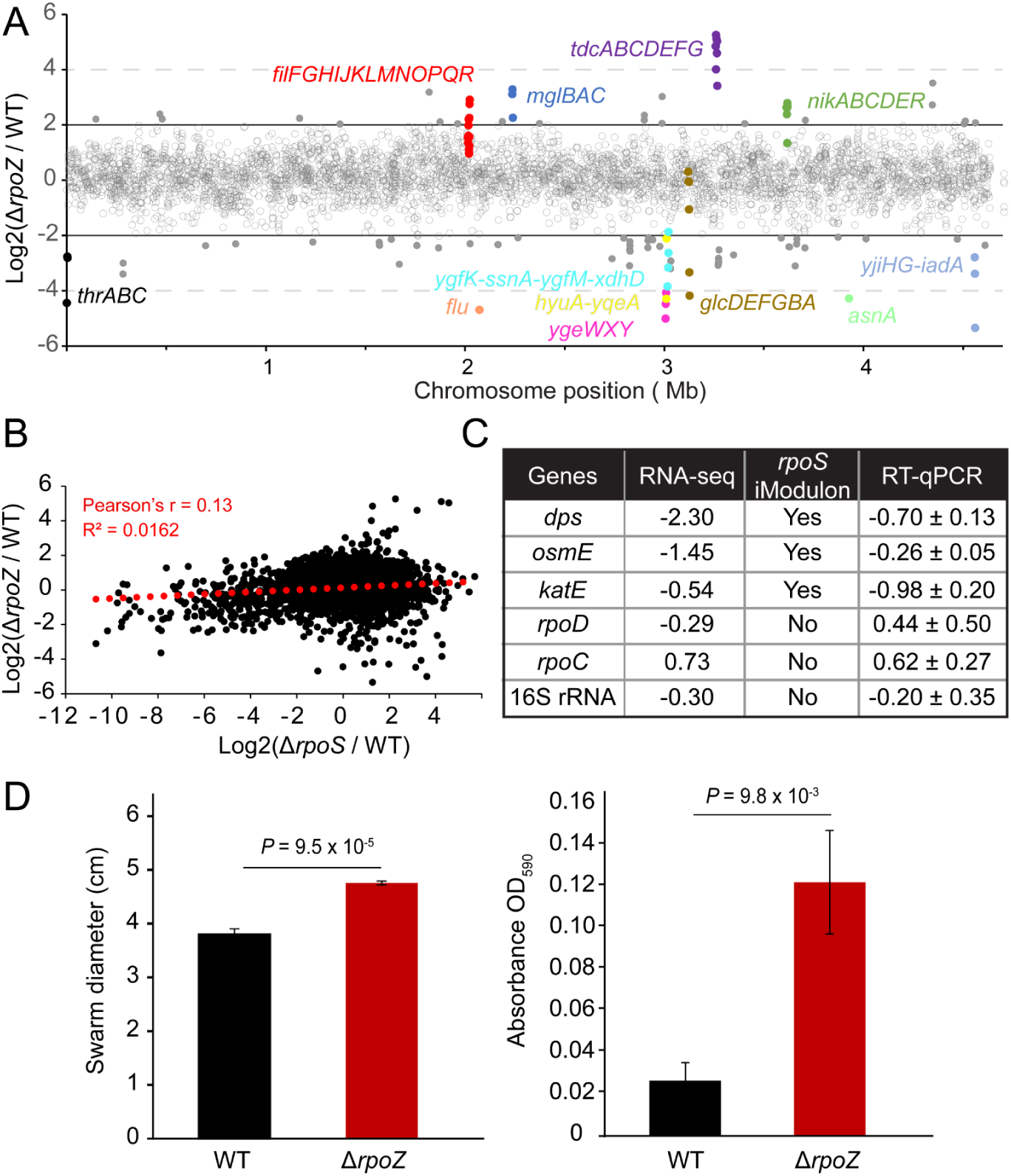
The effects of *rpoZ* deletion on gene expression. (**A**) RNA-seq analysis shows that the expression of all but 108 genes was not significantly affected in the *rpoZ* deletion strain; these genes fall into the ‘gray’ interval log2FC between-2 and +2, indicated with the horizontal black lines. Genes that were up-or downregulated (log2FC >2 or <-2, respectively) are indicated with colored dots above and below these lines. (**B**) RNA-seq shows the lack of correlation between *rpoS* and *rpoZ* deletion datasets. The Pearson correlation and coefficient of determination (R^2^) are shown (**C**) RT qPCR of σ^70^ vs σ^S^ target genes. Data is log_2_(Δ*rpoZ*/WT). Data is shown as mean ± SD (n = 3) for RT-qPCR. (**D**) The deletion of *rpoZ* increases motility (left) and biofilm formation (right). Error bars are SD (n = 4).

In light of this discrepancy, we tested if our Δ*rpoZ* strain has altered biofilm formation and motility, phenotypes commonly associated with the *rpoZ* deletion (*3*). We found that the *ΔrpoZ* strain exhibited increased motility and biofilm formation as compared to the WT strain (**Fig. 2D**). The increased motility is consistent with an increase in expression of the *fliPQR* genes (log₂FC = 2.2 – 2.9), which are required for flagellar biosynthesis. The biofilm formation is a very complex phenotype, and we observed opposite changes in the expression of genes that are known to promote biofilms; *e.g.*, the *flu* (*agn43*) gene is downregulated (**Fig. 2A**), whereas the *bcsA* gene, which is required for cellulose biosynthesis, is upregulated (log₂FC = 1.8).

### CRISPRi screening identifies genes that affect fitness in the ΔrpoZ strain

Our RNA-seq analysis did not reveal any obvious themes in *rpoZ* regulation of gene expression. To identify genetic interactions that affect cell fitness in the absence of ω, we conducted a genome-wide CRISPRi knockdown screen (**Fig. 3A**). In this screen, WT and Δ*rpoZ* strains, harboring an inducible nuclease-dead Cas9 (dCas9) construct, are transformed with a library of guide RNA-expressing plasmids. Because dCas9 binds DNA without cleaving it, it blocks RNAP to effectively silence target genes. Each plasmid in the library targets a specific genomic locus. When dCas9 is induced, robust silencing of these targets creates loss-of-function phenotypes. If silencing a gene provides a growth advantage, the corresponding guide RNAs will be enriched after an outgrowth period; conversely, if silencing hinders growth, those guides will be depleted. By using high-throughput sequencing to compare plasmid abundance before and after outgrowth, we can quantify the fitness effect of each knockdown.

**Fig. 3.**
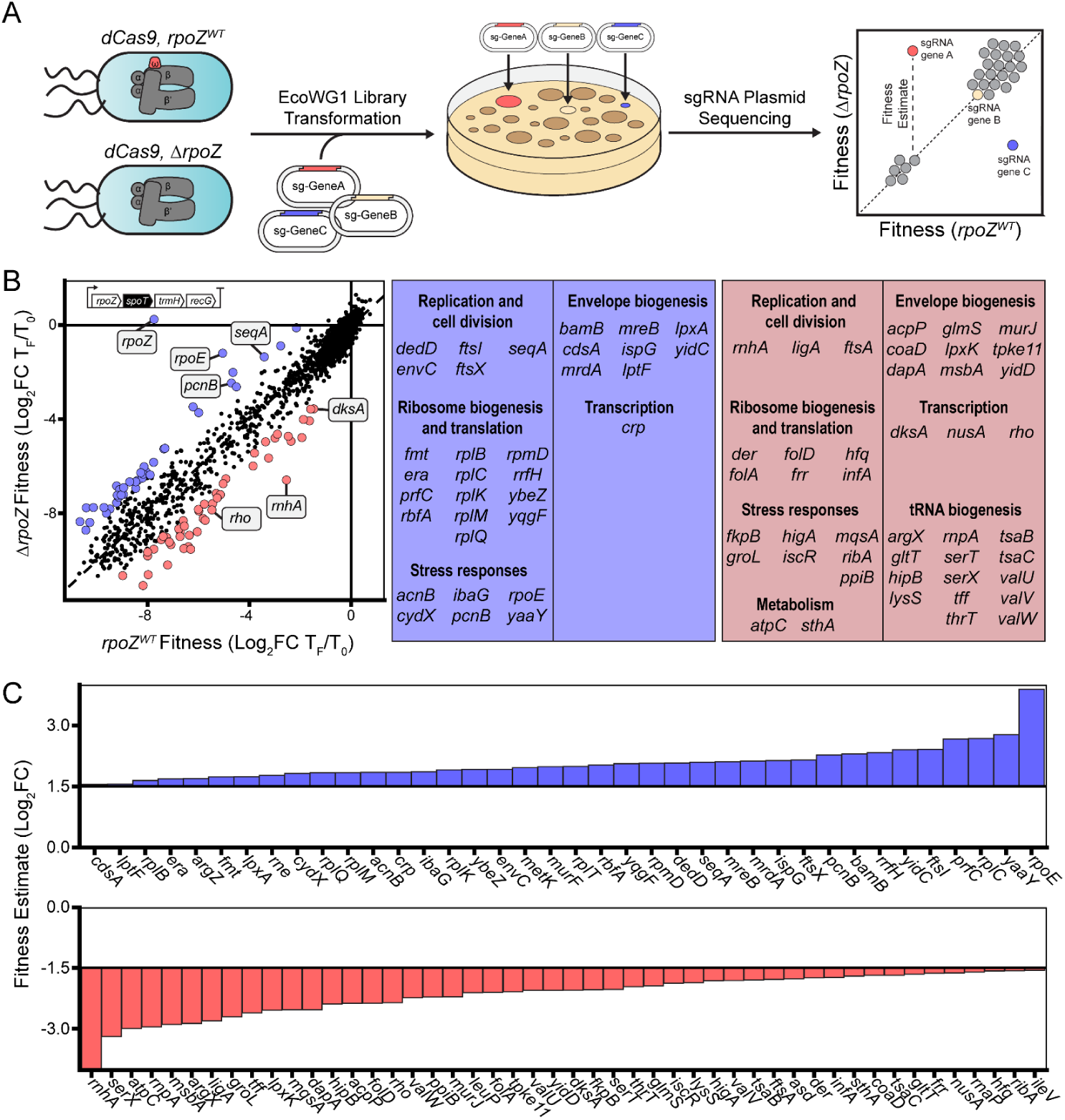
CRISPRi synthetic gene interaction profiling. (**A**) Schematic of the CRISPRi screen. Wild-type (WT) or *ΔrpoZ* cells harboring an inducible *dCas9* construct were transformed with the EcoWG1 whole genome sgRNA library. These pooled libraries were plated on LB media with *dCas9* induction and allowed to grow to saturation for two passages before sgRNA plasmids were isolated from both cell pools. sgRNA plasmid abundance was quantified by Illumina sequencing, and the fold-change in sgRNA abundance over the course of the experiment was quantified and compared across genotypes. (**B**) Comparison of estimated knockdown fitnesses between WT and *ΔrpoZ* cells. Knockdowns that were significantly better tolerated in *ΔrpoZ* cells are labelled in blue and worse tolerated in *ΔrpoZ* cells are labelled in red. The apparent fitness effect of the *rpoZ* sgRNA in *ΔrpoZ* cells is explained by polar effects on the essential *spoT* gene (black). Scores represent the average of four independent screens. Significantly changed genes are grouped into functional categories. (**C**) Estimated significant fitness changes attributed to the *ΔrpoZ* genotype: the knockdowns that are better tolerated are shown at the top, whereas those that are worse tolerated are shown at the bottom.

This analysis identified 77 genes that significantly affect fitness when knocked down (**Fig. 3B**, left). Specifically, 33 knockdowns were better tolerated in *ΔrpoZ* cells (the “UP” group; blue in **Fig. 3B**), while 44 were less tolerated (the “DOWN” group; red in **Fig. 3B**). Proteins involved in translation represent the largest fraction of these hits: ribosome biogenesis factors and ribosomal proteins are prevalent among the UP genes, whereas tRNA synthesis, processing, and modification dominate the DOWN genes. In both groups, envelope biogenesis and stress response functions comprise the second and third largest categories, followed by proteins involved in replication and cell division. Strikingly, only four genes encoding transcriptional regulators were significantly affected: *crp* in the UP group and *dksA*, *nusA*, and *rho* in the DOWN group (**Fig. 3B**, right). While we are not aware of direct links between ω and CRP, a global regulator of transcription initiation, extensive data connect ω to DksA, NusA, and Rho.

DksA and the alarmone (p)ppGpp cooperatively destabilize promoter complexes to reprogram metabolism during stress (*20*). DksA facilitates (p)ppGpp binding within the RNAP secondary channel to mediate most initiation effects, though a secondary (p)ppGpp site at the ω/β’ interface provides DksA-independent regulation (*21*). DksA also inhibits RNA chain elongation (*22*) and is thought to destabilize transcription elongation complexes (TEC) to clear the path for the replisome (*23*). NusA promotes pausing and termination (*2*) and induces folding of the ω C-terminus within the paused TEC (*24*). Rho is an RNA helicase that induces premature termination of antisense, damaged, and poorly translated RNAs to maintain the transcriptome quality and genome stability (*25*). Rho terminates ω-less RNAP more efficiently *in vitro* (*26*), suggesting that the loss of ω may destabilize the TEC. While the lack of strong *ΔrpoZ* phenotypes suggests that any ω-mediated stabilization is likely modest, it may be necessary to support transcription of some operons in which nucleoid-associated proteins act as roadblocks, stalling the elongating RNAP (*27*).

### The lack of ω decreases RNAP processivity

To test whether ω favors processive RNA synthesis, we first compared the ability of the WT and ω-less RNAPs to read through several intrinsic terminators composed of a GC-rich hairpin followed by a U-rich region. We found that ω did not affect termination at P14, T3, and T7 terminators, but decreased RNAP readthrough at λ t_R2_, *rrnB* T1, and *rsxC* terminators (**Fig. 4A**). There was no correlation between the terminator strength and the ω effect: the *rrnB* T1 is the strongest terminator tested here, whereas the *rsxC* is the weakest, requiring NusA for termination (*28*). Interestingly, termination at the ω-sensitive λ t_R2_ and *rrnB* T1 is also affected by DksA and ppGpp (*22*). These results support the proposed stabilization of the TEC by ω and suggest that ω effects are context dependent.

**Fig. 4.**
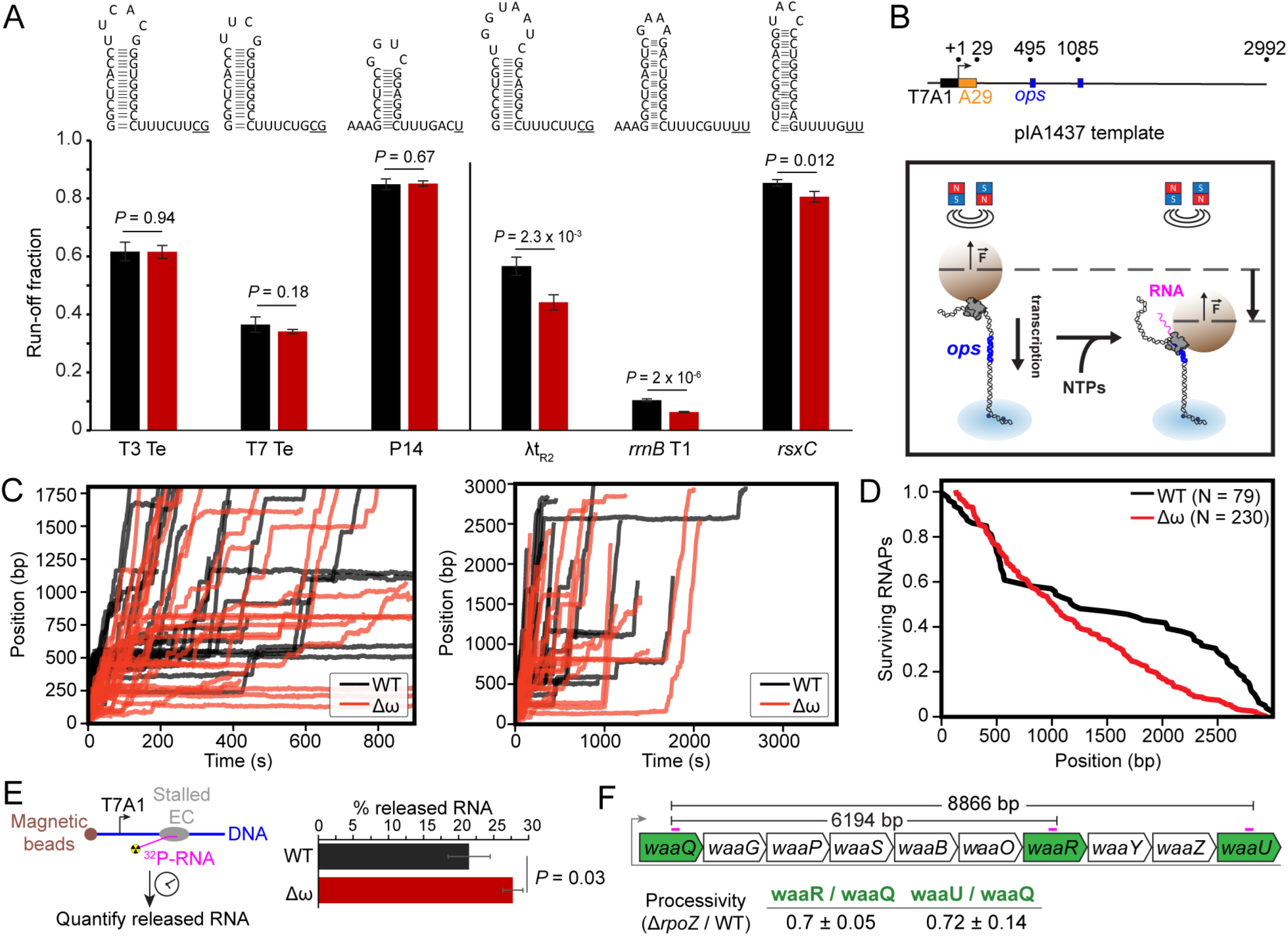
The absence of ω compromises the TEC stability and processivity. (**A**) The absence of ω compromises RNAP readthrough at a subset of intrinsic terminators. WT, black bars. Δω, red, bars. The tested terminators are listed below the bars and the corresponding structures are above. Error bars represent SD (n = 4). A two-tailed t-test assuming unequal variances was used to calculate *P*-values. (**B**) Schematics of the assay in opposing force (OF) configuration, in which a pair of magnets applies force on a paramagnetic bead attached to the *E. coli* RNAP transcribing towards the surface on a tethered DNA template when provided with NTPs. RNAP was stalled in elongation at A29 to establish a stable elongation complex. Location of the *ops* site is indicated. (**C**) WT (black) and Δω (red) RNAP activity traces under 7 pN OF. First 15 min (*left*) and 60 min (*right*) of RNAP elongation traces acquired in the presence of 1 mM NTPs and 37°C; a representative pool of traces (N = 30) of the WT and Δω RNAP is shown. (**D**) Processivity of WT (N = 79) and Δω (N = 232) RNAPs under 7 pN of applied force in OF configuration with 1 mM NTPs at 37°C. (**E**) Lack of ω facilitates the RNA release from the stalled TEC. Error bars represent SD (n = 3). A two-tailed t-test assuming unequal variances was used to calculate *P*-values. (**F**) RT-qPCR shows that *rpoZ* deletion results in a 1.4-fold decrease in processivity in the transcriptionally silenced *waa* operon. RT-qPCR probes are indicated with magenta bars.

To follow the RNA chain elongation over long distances, we carried out high-throughput single-molecule magnetic tweezers transcription experiments on an ∼3 kb DNA template, tracking hundreds of beads in parallel (**Fig. 4B**). Our results reveal that, in a striking contrast with the WT enzyme (**Fig. 4C**), the processivity of ω-less RNAP decays linearly over the long template (3.45 ± 0.44 % every 100 bp, median ± standard deviation of the medians obtained from 100 bootstrap procedures) (**Fig. 4D**), suggesting that the ω-less RNAP consistently enters an unrecoverable stalled state. In contrast, the processivity of the WT RNAP does not show a single rate of decay, with drops corresponding mainly to the early transcribed region and the region where the *ops* site, which induces a very strong RNAP pause (*29*), is located (**Fig. 4CD**). Interestingly, when the ω subunit is absent, the RNAP does not seem to be affected by the presence of the backtrack-inducing *ops* site, both in terms of processivity and of the average time it takes to transcribe the *ops*-containing region (**Fig. S4**).

To determine if the ω-less RNAP forms unstable TECs, we immobilized halted TECs, in which the nascent RNA is isotopically labeled, challenged them with competitors, and measured the release of the nascent RNA (**Fig. 4E**). We found that after 30 min incubation, 21.5% of RNA was released from the WT TECs, as compared to 27.7% of RNA released by the ω-less RNAP.

We next probed the effects of ω on RNAP processivity *in vivo* in the *waa* operon (**Fig. 4F**). The *waa* operon is horizontally acquired and is known to be silenced by Rho and H-NS (*30*). A decrease in RNAP processivity would lead to the reduced expression of the downstream genes, *waaR* and *waaU*, relative to the first gene, *waaQ*. We used RT-qPCR assay to determine the *waaR*/*waaQ* or *waaU*/*waaQ* ratios in RNAs isolated from the WT and Δ*rpoZ* strains. We found that the *rpoZ* deletion reduced processivity to 70% of that in the WT strain (**Fig. 4F**).

Collectively, these results demonstrate that the loss of ω increases readthrough of a subset of intrinsic termination signals, reduces stability of halted TECs, and compromises processivity *in vitro* and *in vivo* (**Fig. 4**). While these differences are modest, consistent with the mild phenotype of Δ*rpoZ* strains, they likely amplify under specific cellular stresses.

### Deletion of rpoZ suppresses the sensitivity of genome stability mutants to DNA-damaging agents

Our data support a model in which ω-less RNAP is more prone to dissociation. While the TEC destabilization might compromise the efficient synthesis of long RNAs, it likely promotes TRC resolution and transcription-coupled repair, two essential processes for maintaining genome stability. A stalled RNAP can hinder repair enzymes’ access to damaged DNA and block the replisome path. NusA can promote repair (*31*), whereas DksA and Rho both facilitate TRC resolution (*32, 33*) and confer partial resistance in *E. coli* to agents inducing double-stranded DNA breaks (DSBs) (*34–36*).

Although our CRISPRi screen did not identify specific repair factors, we found that several genes controlling DNA replication (*seqA*) and cell division (*dedD, envC, ftsI*, and *ftsX*) became less critical in the *ΔrpoZ* background (**Fig. 3B**). We posit that the loss of ω mimics the protective effects of DksA and Rho; consequently, the deletion of *rpoZ* should improve the growth of DksA-or Rho-defective strains in the presence of genotoxic compounds.

To test this idea, we constructed strains with the Δ*dksA* or a *rho*-down allele, an IS2 insertion in the *rho* leader region that reduces Rho cellular concentration 2.5-fold (*37*), carrying either the WT or Δ*rpoZ*. The double deletion strains had modest growth defects in liquid culture (**Fig. S3**) but grew better than the Δ*dksA* and *rhoL::ΩIS2* strains on subinhibitory concentrations of mitomycin C (MMC), phleomycin (PHL), and nalidixic acid (NAL), an effect that was particularly pronounced with the *rho* allele (**Fig. 5**).

**Fig. 5.**
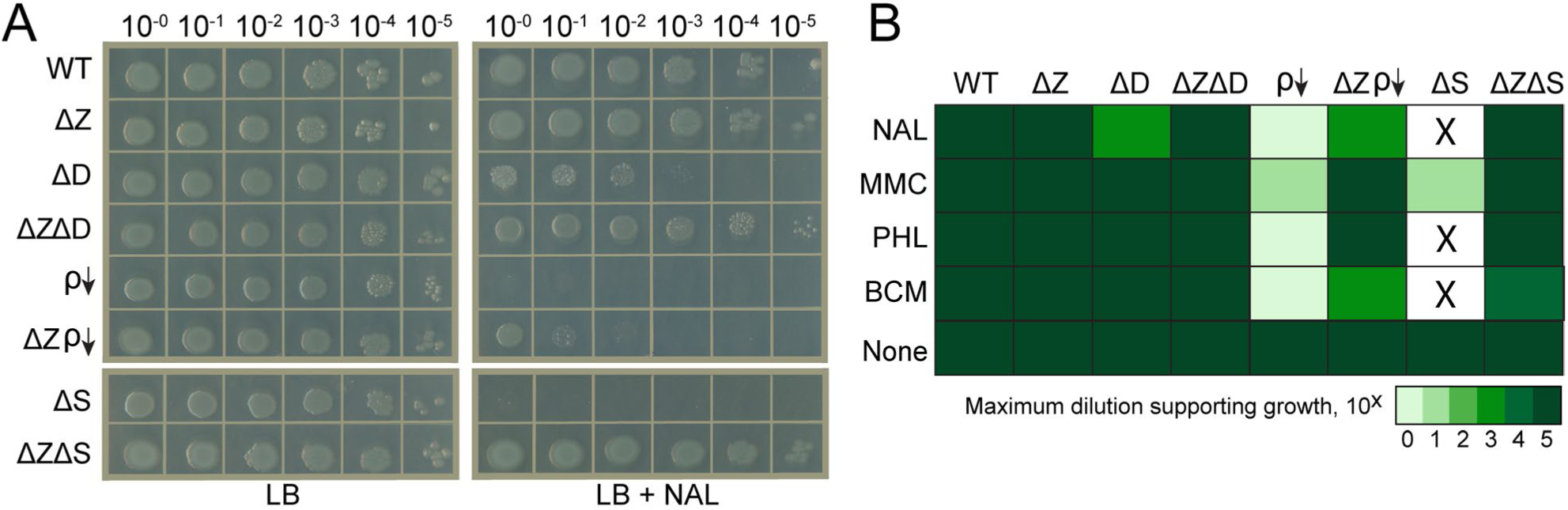
Deletion of *rpoZ* (ΔZ) suppresses the sensitivity of Δ*dksA* (ΔD), *rho* knockdown (*rhoL*::IS2, ρ↓) and Δ*seqA* (ΔS) strains to DNA-damaging agents. (**A**) Representative plates showing that ΔZ suppresses the genotoxic effects of NAL. (**B**) Heatmap showing the effects of different DNA-damaging agents on tested strains. 2 μg/ml NAL, 0.25 μg/ml MMC, 1 μg/ml PHL, and 20 μg/ml BCM were used; see **Fig. S5** for images of plates supplemented with BCM, MMC, and PHL.

These agents promote DSBs by different mechanisms: the prodrug MMC, once activated, induces interstrand crosslinks of guanine residues; PHL catalyzes free radical-mediated DNA breaks; and NAL targets DNA gyrase to trap its covalent DNA-bound intermediates.

TRCs are unavoidable since the replisome moves ∼25 times faster than RNAP and will run into the transcribing RNAP, particularly on heavily transcribed genes such as those that encode ribosomal RNAs (*38*). To minimize these conflicts, *E. coli* cells carefully time the initiation of replication (*39*). The SeqA protein binds to hemimethylated GATC at *oriC* and blocks premature recruitment of DnaA to ensure that the chromosome is replicated only once per cycle. In Δ*seqA* cells, origins fire repeatedly and asynchronously (*40*), increasing the TRC frequency. It follows that under these conditions, the loss of ω should have a protective effect; consistently, the *seqA* knockdown is better tolerated in the Δ*rpoZ* background (**Fig. 3B**). To confirm this connection, we compared the sensitivity of the Δ*seqA* strain to DNA damage-inducing compounds with and without *rpoZ* (**Fig. 5**). We found that the Δ*seqA* strain failed to grow on NAL, PHL, and BCM, and barely grew on MMC; strikingly, Δ*rpoZ* restored viability of the double deletion strain (**Figs. 5** and **S5**). We also tested the sensitivity of Δ*envC* and Δ*rnhA* strains to genotoxic stress and the presence of *rpoZ* (**Fig. S6**). Our results show that the deletion of *envC* confers sensitivity to BCM and PHL, which is suppressed by Δ*rpoZ.* By contrast, the Δ*rnhA* strain was largely insensitive to DNA-damaging agents and *rpoZ* (**Fig. S6**).

Curiously, while we show that Δ*rpoZ* suppresses defects of *dksA*, *rho*, and *seqA* during genotoxic stress (**Fig. 5**), our CRISPRi screen categorized these genes differently (**Fig. 3**); interference with *dksA* and *rho* expression was expected to reduce the fitness of the Δ*rpoZ* strain. Consistent with this, we found that the deletion of *rpoZ* compromised the growth of Δ*dksA* and *rho*-down mutants but had little effect on Δ*seqA* under optimal conditions (LB at 37°C; **Fig. S3**). This discrepancy is likely explained by differing growth requirements: while Δ*rpoZ* dramatically improves growth under genotoxic stress, it is neutral or even detrimental during optimal growth in LB, conditions used both for CRISPRi and growth assays.

## Discussion

Here, we reexamine the role of the smallest RNAP subunit in gene expression of *E. coli.* Surprisingly, we did not observe activation of the σ^S^ regulon upon the *rpoZ* deletion (**Fig. 2**) or changes in plasmid DNA topology (**Fig. S2**), a discrepancy that we attribute to likely polar effects caused by the *rpoZ* disruption in earlier studies (*7, 12*). Instead, our past and present results demonstrate that the loss of ω sensitizes RNAP to termination triggered by Rho (*26*) and some intrinsic hairpin-dependent signals (**Fig. 4A**), reduces TEC stability and processivity *in vitro* (**Fig. 4DE**), and augments Rho-mediated polarity *in vivo* (**Fig. 4F**). Our CRISPRi screen suggested that the loss of ω may promote survival under conditions when RNAP and replisome collide frequently, *e.g.,* when *seqA* silencing leads to runaway replication (**Fig. 3**). Indeed, we found that Δ*rpoZ* fully suppressed the dramatic growth defects of Δ*seqA* and promoted survival of strains lacking functional Rho and DksA in the presence of DNA damaging agents (**Fig. 5**). We hypothesize that ω/RPB6 homologs maintain genome integrity across all domains of life by balancing processive long-RNA synthesis with the timely release of RNAP to facilitate TRC resolution.

Cellular RNAPs are obligately processive: once an enzyme clears the promoter, it must synthesize the entire RNA transcript in a single, uninterrupted event. For long operons, this feat can span a duration nearly equal to a cell cycle. To prevent premature release, the TEC is inherently stable, sometimes recruiting accessory antitermination factors to support transcription of challenging operons (*41*). In this scenario, the ω ‘latching’ action supports the TEC processivity (**Fig. 6A**).

**Fig. 6.**
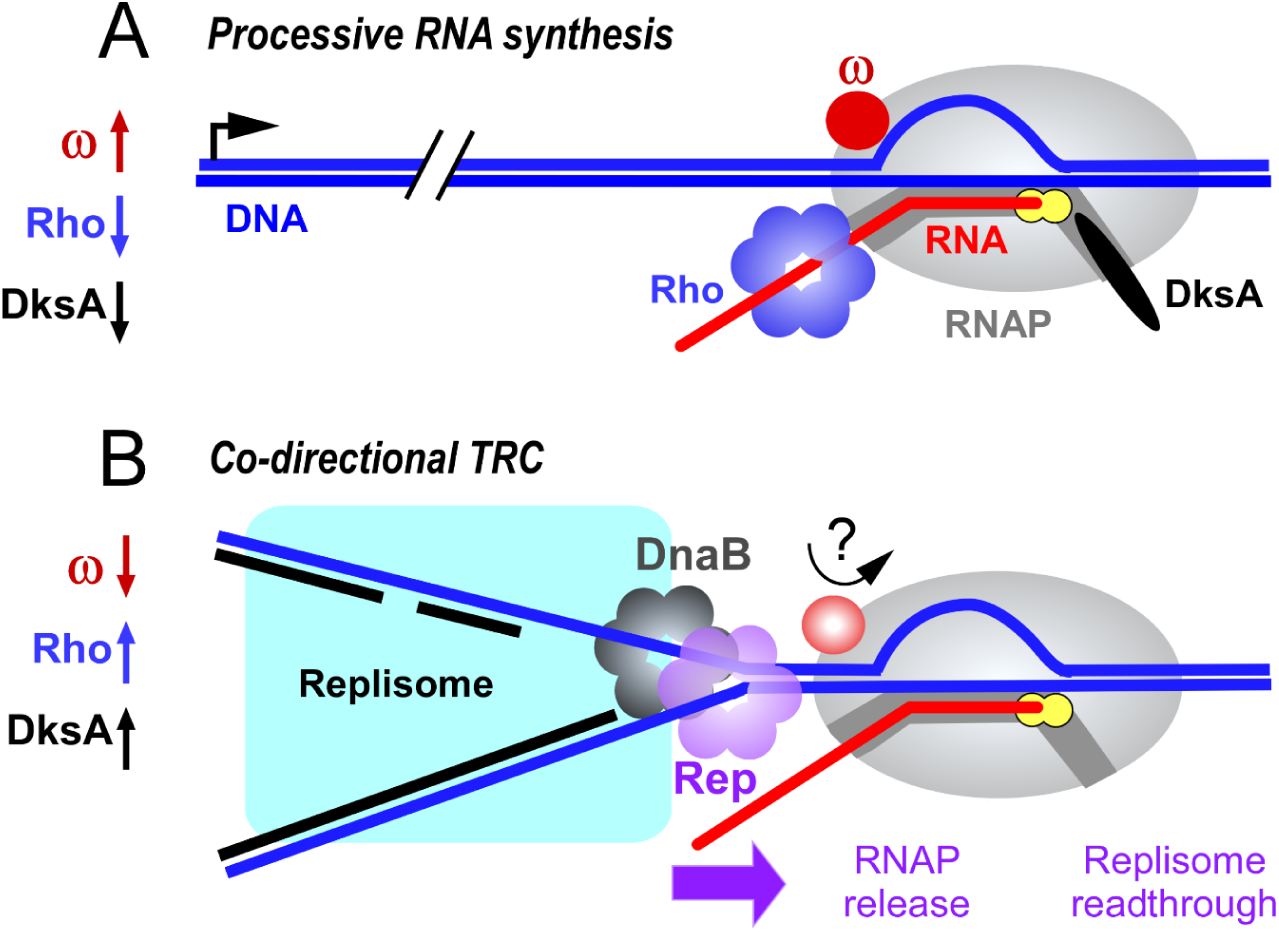
Context-dependent effects of ω, Rho and DksA. (**A**) ω promotes while DksA and Rho inhibit synthesis of long RNAs. (**B**) During co-directional TRC, ω release (possibly potentiated by Rep or another component of the replisome), DksA and Rho destabilize the TEC to enable replisome readthrough.

However, an overly stable TEC becomes a liability when it blocks unimpeded passage of the replisome or the DNA repair machinery’s access to a site of DNA damage (**Fig. 6B**). The fast-moving *E. coli* replisome (1000 bp/s) repeatedly encounters slow TECs (40 – 80 bp/s) in either co-directional or head-on orientation. Here, we focus on co-directional collisions, which are less disruptive (*38*) but far more frequent: during rapid growth, the seven *rrn* operons, aligned with the direction of replication, initiate RNA synthesis every second (*42*). The mild consequences of co-directional TRCs may stem from at least two mechanisms. The 3’-5’ helicase Rep associates with the replisome through direct contacts with DnaB, loads onto the single-stranded DNA, and dislodges the RNAP ahead of the fork (*43*). The Rep helicase action is likely sufficient to disrupt the minimal contacts between RNAP and the upstream DNA duplex (*44*). Additionally, the specialized TECs that transcribe *rrn* operons (*45*) are resistant to pausing/backtracking and transcribe twice as fast as their unmodified counterparts (*46*); this pause-free state may be critical for facile collision resolution, as RNAP stalling is known to exacerbate the severity of TRCs (*47*).

TRC resolution is aided by transcription factors that destabilize the TEC, such as DksA and Rho (*32, 36*), by substitutions in RNAP (*48*), or by (p)ppGpp (*49*). Except for Rho, the molecular details of these destabilizing mechanisms remain unclear. Similarly, it remains to be determined how the loss of ω makes the TEC less stable; the original ‘latching’ model was based on the *Thermus aquaticus* RNAP structure (*4*) in which ω contacts with the β and β’ subunits are more extensive. In *E. coli*, ω appears to be loosely bound to the α_2_ββ’ core and readily dissociates during purification (*50*). This instability is further evidenced by its selective degradation in the stationary phase (*51*) and its displacement by the translating ribosome that collides with a stalled RNAP (*52*). An analogous mechanism may exist in *B. subtilis,* where the ATPase HelD induces dramatic conformational changes that facilitate RNAP recycling and promote ω dissociation (*53*). It is thus possible that, in addition to the helicase action of Rep, the replisome collision with the TEC may also trigger ω release. The precise mechanism by which the ω subunit stabilizes the TEC is the subject of ongoing study.

Transcription-coupled DNA repair is another essential pathway that maintains genome integrity. Through pervasive transcription, RNAP scans the DNA and stalls at various lesions, marking them for repair (*54*). While the stalled RNAP transiently masks the lesion, forward translocation induced by Mfd (*55*) or backtracking induced by the UvrD helicase and ppGpp (*56–58*) exposes the lesion to the UvrA_2_B complex, priming it for repair. Neither *mfd* nor *uvrD* has been identified in our CRISPRi screen (**Fig. 3**), and it remains to be determined if ω influences either repair pathway; notably, however, the ppGpp-binding site implicated in DNA repair is located at the ω/β’ interface (*58*). The loss of ω (as well as the destabilizing action of DksA/Rho) may also inhibit repair by promoting the premature release of lesion-stalled RNAP, delaying repair until subsequent rounds of transcription.

The story of ω is one of perpetual redefinition: initially dismissed as a contaminant (*50*), ω was recognized as a universal RNAP assembly chaperone (*4*), and may act as a genome guardian through ppGpp-mediated DNA repair (*58*). Our findings suggest a novel role for ω in resolving the lethal collisions between transcription and replication machinery. We propose that ω and its universal homologs serve a fundamental purpose by fine-tuning TEC stability during replication and repair to safeguard the genome against fatal molecular roadblocks.

## Materials and methods

### Bacterial strains and plasmids

*E. coli* MG1655 strains, plasmids, and oligonucleotides used in this study are listed in **Table S2**. *E. coli* strain with the *rpoZ* clean deletion was constructed by using the λ Red recombineering method and CRISPR-Cas9 counterselection (*17*). Kanamycin resistance marker was flipped out by pCP20 (*59*). All other strains were generated by P1 transduction of appropriate P1 lysate either in IA900 (for generating single gene deletion mutants) or IA909 (for generating double gene deletion mutants). The resulting deletion strains were confirmed by PCR with a forward primer in the *kan^R^* gene (*neo*_F) and matching reverse primers in the target gene.

### E. coli MG1655 ΔrpoZ clean deletion strain construction

Donor DNA oligonucleotide (80 bp long) used for recombination was designed to target the lagging strand of replicating DNA with upstream and downstream homology regions spanning the *rpoZ* gene. The donor DNA was 5’ end-modified with four phosphorothioated bases to prevent degradation by host endonucleases. pBK1, pKDsg derivative, used for counterselection, was constructed by p453 (pKDsg) digestion with BsmBI-v2 (NEB) and consequent ligation with synthetically designed (Millipore Sigma) annealed 24-bp primer duplex using T4 DNA ligase (NEB). Primers contained the overlapping 20 bp protospacer region, binding adjacent to the corresponding PAM (5’-NGG) sites in the *rpoZ* chromosomal sequence. The sequence of the pBK1 construct was confirmed by Nanopore sequencing (Plasmidsaurus).

The IA765 strain was first transformed with p323 (pCas9cr4) plasmid, selected on LB with 25 mg/l chloramphenicol (Cm), and subsequently transformed with the single guide RNA encoding plasmid (pBK1). Cells possessing both plasmids were grown in SOB medium with spectinomycin (Sp, 50 mg/l) and 25 mg/l Cm at 30°C. At OD_600_ ≈ 0.5, λ Red was induced with 0.2% (w/v) L-arabinose, and cells were grown for another 15 min. Cells were chilled on ice for 5 min and washed with 10% cold glycerol before freezing at-80°C while in 10 % glycerol. The next day, donor DNA for recombineering spanning the *rpoZ* sequence was electroporated into the cells using standard Ec1 parameters. After an hour recovery at 30°C in 1 ml SOC, the cells were plated on LB medium supplemented with Sp (50 mg/l), Cm (25 mg/l), and anhydrotetracycline (aTc, 100 μg/l) and incubated overnight at 30 °C to select for survivors of the CRISPR–Cas9 selection. Resulting colonies were screened with specific primers to confirm the chromosomal *rpoZ* gene deletion.

To eliminate the pBK1 plasmid, cells were inoculated in LB medium and grown at 37°C overnight, and then plated on LB. Individual colonies were patched on LB and LB supplemented with 50 mg/l Sp to assess the loss of Sp resistance. The p323 (pCas9cr4) plasmid was cured by the p454 (pKDsg-15a) plasmid. Upon transformation p454 (pKDsg-15a) into *E. coli* MG1655 *ΔrpoZ* strain, still containing pCas9cr4 plasmid, cells were recovered in SOC medium and plated on LB supplemented with Sp (50 mg/l), Cm (25 mg/l), and aTc (100 μg/l) overnight for selection. The next day, colonies were patched to assess the loss of Cm resistance. The p454 (pKDsg-15a) plasmid was cured by growth at 37°C. The final strain was sequenced by the IDI-GEMS at The Ohio State University.

### RpoZ distribution analysis

We searched the ω InterPro entry (IPR003716, bacterial RNAP subunit omega) in AnnoTree version r214 (*60*) with default settings to detect the distribution of ω homologs across the phylogenetic tree of Bacteria. The phyla that appear to largely lack ω homologs were manually checked using NCBI BLASTp (https://blast.ncbi.nlm.nih.gov/Blast.cgi) with default settings, using ω homologs from the same phylum as the query. The ω homolog structures were predicted by AlphaFold3 (*61*) and visualized with PyMOL 3.0.3.

### Purification of Δω RNAP

*E. coli* BL21 derivative that carries the chromosomal Δ*rpoZ* allele (IA356) was constructed by P1 transduction of the *rpoZ::Kan^R^* allele from the Keio collection(*62*), transformed with the pIA299 plasmid that expresses *rpoA, rpoB*, and *rpoC* genes, and plated on LB plates supplemented with 100 mg/l carbenicillin (Carb) for selection. On day 2, colonies were inoculated in 12 ml LB supplemented with 100 mg/l Carb and grown at 37°C overnight. On day 3, culture was diluted 1:100 into 600 ml TB broth (supplemented with 100 mg/l Carb) and grown to OD_600_= 0.6 – 0.7 at 37°C, at which point culture was induced by the addition of 0.5 mM IPTG for 2 – 3 h at 37°C.

Culture was harvested by centrifugation (6,000×*g*, 10 min, 4°C) and lysed by sonication in 40 ml lysis buffer (500 mM NaCl, 50 mM Tris-HCl, pH 6.9, 5% glycerol, 0.2 mM β-ME) supplemented with 1× Complete EDTA-free protease inhibitors cocktail (Roche), and 1 mg/ml lysozyme. Lysate was cleared by centrifugation at 20,000×*g* for 30 minutes at 4°C. Cleared lysate was loaded on Ni^2+^-NTA column (Cytiva) equilibrated in lysis buffer. Δω RNAP was washed with lysis buffer supplemented with 10 mM and 20 mM Imidazole, before eluting with Ni-NTA Elution buffer: (20% lysis buffer/80% HepA buffer (50 mM Tris-HCl, pH 6.9, 5% glycerol, 1 mM β-ME) with 300 mM Imidazole.

Eluted Δω RNAP was diluted (2×) with HepA buffer to lower salt content and loaded onto a 1 ml Heparin HiTrap column equilibrated in HepA buffer. The sample was washed with 5 column volumes of HepA buffer before eluting in a linear gradient of 0 – 1.5 M NaCl in HepA. Peak fractions at around ∼40 mS/cm were collected. The eluted protein was diluted (3×) with HepA buffer and loaded onto Resource Q (Cytiva) column equilibrated in HepA buffer. Washing and elution steps were the same as for Heparin column, except peak fraction of RNAP elution was collected at ∼24 mS/cm.

Eluted Δω RNAP was diluted (4×) in HepA buffer and concentrated by centrifugation at 4200 rpm at 4°C, in a concentrator tube (Vivaspin 100K MWCO). Purified and concentrated Δω RNAP was mixed with glycerol (50% final), aliquoted, and stored at-80°C.

### In vitro transcription termination assays

The WT core RNAP and σ^70^ transcription initiation factor were purified as described in (*63*). RNAP holoenzymes were assembled by mixing the core RNAP with a 3-fold molar excess of σ^70^, followed by incubation at 37°C for 15 min. DNA templates were generated by PCR amplification with primers listed in **Table S2** and purified by QIAquick PCR Purification Kit (Qiagen). The transcription template was prepared by PCR amplification from pIA263 (T7 termination signal), pIA265 (P14 termination signal), pIA692 (*rrnB* T1 termination signal), and pIA1239 (NusA-dependent *rsxC* termination signal). Halted complexes (HCs) were formed at 37°C for 15min by CTP deprivation after mixing 60 nM RNAP holoenzyme (WT and Δω) with 30 nM DNA template, 100 μM 5’-ApU, start (2 μM ATP, 10 μM GTP, 10 μM UTP), and 0.1 μCi/μl [α-^32^P]-ATP in TGA^2^ buffer. After the addition of 2 μM NusA (for the *rsxC* terminator only), the reactions were incubated for 2 min at 37°C. Transcription was restarted by addition of a pre-warmed (at 37°C) chase (200 μM each NTP and rifampicin 20 μg/ml) and incubated at 37°C for 20 min. After 20 min, reactions were quenched by the addition of an equal volume of stop buffer (45 mM Tris-borate, 10 M urea, 20 mM EDTA, 0.2% xylene cyanol, 0.2% bromophenol blue, pH 8.3) and separated by 6 – 8% denaturing PAGE (7 M urea, 0.5X TBE).

Samples were heated for 2 min at 95°C and separated by electrophoresis in denaturing acrylamide (19:1) gels (7 M urea, 0.5 × TBE) of various concentrations (6 – 12%). RNA products were visualized and quantified using PhosphorImager Typhoon 9000 FLA (GE Healthcare), Image Quant v5.2, and Microsoft Excel.

### Elongation complex stability assay

Biotinylated pIA447 DNA template was prepared by PCR amplification and purified using QIAquick PCR Purification Kit. In an elongation stability assay, 3 μl of Dynabeads Streptavidin T1 magnetic beads (ThermoFisher, cat# 65601) were washed once with 200 μl of PBS, and then twice with 100 μl of Coupling Buffer (CB: 20 mM Tris-HCl, pH 8.0, 1 mM EDTA, 0.5 M NaCl). After washing, the beads were resuspended in 20 µl CB containing 160 nM biotinylated pIA447 DNA template. The bead suspensions were incubated for 30 min on a rotary shaker (1000 rpm) at 25°C. After incubation, the beads-DNA complex was washed twice with 200 μl TGA2 (20 mM Tris-acetate, 20 mM Na-acetate, 2 mM Mg-acetate, 5% glycerol, 5 mM β-ME, 0.1 mM EDTA, pH 7.9) + 0.1 mg/ml BSA. The washed beads-DNA was mixed into a 55 μl reaction containing 30 nM RNAP holoenzyme (WT/Δω), 100 μM ApU, start NTPs (1 μM ATP, 5 μM GTP, and 5 μM CTP), 0.1 μCi/μl [α-^32^P]-ATP, and 500 nM GreA in TGA2. Reactions were incubated for 18 min at 37°C. The beads were washed 3 times with 200 μl of TGA2 + 0.1 mg/ml BSA. The resulting beads were resuspended in 20 μl reaction containing 0.4 M NaCl, 0.2 mg/ml heparin, and 1 μM T7A1 DNA (**Table S2**). The heparin and T7A1 DNA were used as competitors to prevent RNAP rebinding to the released template DNA. After 30 min on a rotary shaker (1000 rpm) at 25°C, the tube was held against a magnetic rack to separate supernatant and beads. The supernatant was mixed with an equal volume of 2xSTOP buffer (10 M urea, 60 mM EDTA, 45 mM Tris-borate, pH 8.3, 0.1% bromophenol blue, and 0.1% xylene cyanol). The beads were mixed with 20 μl TGA2 + 20 μl 2×STOP buffer. The samples were heated for 3 min at 95°C, and resolved in a 10% acrylamide (19:1) gel (7 M urea, 0.5×TBE). The gels were scanned with Amersham Typhoon 5 and quantified by Fiji v2.17.0.

### RNA sequencing

Single colonies of strains IA765 and IA1003 were inoculated overnight in 2.5 ml of LB at 37°C. Overnight cultures were diluted 1:100 in 5 ml LB and incubated at 37°C until the OD600 of 1.2. After lysing 1 ml cell pellet with 1 ml RNAzol RT (Sigma) and adding 0.4 ml RNase-free water, 7× volume excess of buffer RLT. After that, the standard protocol for Qiagen RNeasy Mini was followed. Fraction of isolated RNA (2500 ng) was further treated with 15 units of DNase I and RNase inhibitor in DNase I buffer for 30 min at 37°C to remove any remaining DNA and ensure RNA integrity. Samples were cleaned up by Monarch Spin RNA Cleanup Kit (NEB) and submitted for sequencing at IDI-GEMS at The Ohio State University.

Adapter and low-quality reads were trimmed using TrimGalore v0.6.7 (*64*) in paired-end mode and one base was trimmed from the 5’ end of all reads in two-colors mode (parameters: --paired --trim-n - clip_R1 1 --clip_R2 1 --2colour). Reads were aligned to *E. coli* MG1655 genome (ASM584v2) using bowtie2 v2.4.4 (*65*) with very sensitive, end-to-end alignment, and dovetailed alignments allowed (Parameters: --very-sensitive --no-mixed --no-discordant --dovetail-X 1000). The alignment files were converted to BAM format and indexed by SAMtools v1.23 (*66*). The alignment files were subjected to quantification using featureCounts v2.0.3 (*67*) (parameters: --countReadPairs --minOverlap 15). Counts per million (CPM) was calculated in Excel.

### Reverse transcriptase quantitative PCR (RT-qPCR)

Strains were grown in LB at 37°C to an OD_600_ of 1.2, and the cells were mixed with 0.2 volume of ice-cold growth stop solution (5% phenol and 95% ethanol) to inactivate cellular RNase before being collected by centrifugation at 8,000×*g* for 3 min. Total RNA from the aqueous phase of RNAzol RT extracted sample was isolated by Monarch Total RNA miniprep kit (NEB; cat# T2010). RNA samples were treated with TURBO DNase (ThermoFisher, cat# AM2239) before RT-qPCR. A total of 100 ng RNA samples were used with iTaq Universal SYBR green one-step kit (Bio-Rad, cat#1725150) and analyzed on a CFX96 system (Bio-Rad). Samples without reverse transcriptase were used as a negative control to ensure the absence of DNA contamination. The quantification cycle (Cq) values were calculated using CFX Manager v3.0 in regression mode. The gene expression level was analyzed by the threshold cycle (2^−ΔΔCT^) method; the *ihfB* gene was used as a reference.

### High-resolution electrophoresis analysis of plasmid topology

Cells were grown in LB to an OD of 0.4 and collected for plasmid preparation using Monarch Plasmid Miniprep Kit (NEB). 500 ng plasmid was resolved in 1% agarose gel (SeaKem LE Agarose; VWR, cat#12001-868) containing 5 μg/ml chloroquine. Electrophoresis was performed at 27 V (2 V/cm) in 1×TAE buffer for 17 hours at 4°C. The gel was washed for 15 min with water three times before being stained with SYBR Gold. The gel was scanned with Amersham Typhoon 5 and quantified by Fiji v2.17.0. A rolling ball background subtraction was performed before plotting in Fiji.

### Motility assays

To examine motility, single colonies of WT (765) and *ΔrpoZ* (1003) clean deletion strains were stabbed in the center of a motility agar plate (LB containing 0.3% agar) followed by an incubation at 30 °C for 2 days. Strain motility was measured by measuring swarm motility diameter at 24 h and 49 h time points.

### Biofilm formation assays

WT (765) and *ΔrpoZ* (1003) strains were grown at 37°C in LB Lennox media (250 rpm) to prepare an overnight starting culture. The next day, 1 ml of the starting cultures was spun down and resuspended in LB Lennox to adjust the OD_600_ = 3 before diluting them 1:100 at a final OD_600_ = 0.03 in 200 μl LB Lennox. Strains were left for incubation at 25°C for 5 days under static conditions in Costar 96 well plate (flat bottom, non-treated, polystyrene) to allow biofilm formation. The plate was washed 3 times with deionized (DI) water, stained by the addition of 0.1% crystal violet dye for 20 min at room temperature, and washed 3 times again with DI water. 10% acetic acid was added to the wells and incubated for 20 min at room temperature with gentle swirling. Finally, solutions were transferred to a fresh 96-well plate, and OD_600_ was measured at 590 nm with the Epoch 2 plate reader (BioTek). Data was exported and processed using Microsoft Excel.

### General growth assays

MG1655 derivative double and single deletion strains (indicated in the figure legend) were grown at 37°C in LB Miller media (250 rpm) to prepare an overnight starting culture. The next day, overnight cultures were spun down and resuspended in LB Miller to a final OD_600_ = 0.03 in 200 μl LB Miller. Strains were grown at 37°C for 24 hours in the Epoch 2 plate reader (BioTek) using Greiner 96 Cellstar plate.

OD_600_ readings were taken every 15 min, with continuous shaking at double orbital. Data was exported and processed using Microsoft Excel.

### CRISPRi screening

The assay was carried out using the EcoWG1 library (*68*), following the protocol described in the previous study (*69*). Two independent, clonal cultures from each line were transformed, via electroporation, with the EcoWG1 sgRNA library to a minimum library coverage of 100× (2 × 10^6^ transformants), recovered, and stored as-80°C glycerol stocks. These stocks were then thawed and used to seed 50 ml LB + 20 µg/ml kanamycin cultures. These cultures were grown overnight, to saturation and without dCas9 induction, to a titer of ∼6 × 10^9^ (OD_600_ = 6.0) viable cells per ml. Each culture was diluted 1:500 in MinA +1 mM MgSO_4_ to 12 × 10^6^ cells/ml (OD_600_ = 0.012) and 83.5 µl of culture was spread on three MinA +20 µg/ml kanamycin, 200 ng/ml anhydrotetracycline plates supplemented with 0.2% of the indicated carbon source. The plates were incubated at 37°C for 24 hours, to saturation. The plates were then manually scraped with 3 ml PBS, generally obtaining 1 ml of∼1.2 × 10^10^ viable cells per ml (OD_600_ = 120; OD_600_ of 1:100 dilution = 1.2). This mix was diluted 1:10,000 in PBS (1:100 then 1:100) to 12 × 10^6^ cells/ml, and 83.5 µl of culture was spread on three more MinA + 20 µg/ml kanamycin, 200 ng/ml anhydrotetracycline plates supplemented with 0.2% of the indicated carbon source. The plates were incubated for another 24 hours at 37°C, to saturation. The plates were again scraped with 3 ml PBS and 50 µl (equivalent of OD_600_ = 6.0) of the resulting slurry was then miniprepped to isolate plasmid.

The sequencing library preparation is composed of two sequential PCRs of the input sgRNA plasmids, which add the sequencing adapters and indexes. The first PCR amplifies from the sgRNA promoter region to a region ∼300 bp downstream of the sgRNA. The primer overhangs add the read 1 primer binding site, the index 2 (i5) sequence, the index 2 primer binding site, and the index 1 (i7) primer binding site. The second PCR adds the flow cell attachment sequences, P5 and P7, as well as the index 1 (i7) sequence. This creates a 487 bp final amplicon.

For the library preparation, 100 ng of sgRNA plasmid is added to the first PCR with the appropriate PCR 1 forward primer (index 2) and a general reverse primer, in a hot-start Phusion PCR. This reaction is subjected to 9 PCR cycles. 5 μl of this 50 μl reaction is added to the second PCR with a general forward primer and the appropriate PCR 2 reverse primer (index 1), in another hot-start Phusion PCR. This reaction is subjected to another 9 PCR cycles. The final product is purified with 1× Ampure XP beads, and the total product is quantified. The final pool of plasmids is pooled according to the desired ratio of reads per sample and run on a 1% agarose gel. If apparent primer dimers are present, the library is subjected to another 1× Ampure XP cleanup, before diluting the library to approximately 1.192 ng/μl final concentration. The library molarity is then quantified using a Colibri library quantification kit and adjusted to a 2 nM final concentration.

For sequencing, the library is diluted and denatured according to standard Illumina protocols, alongside custom HPLC-purified read 1, index 1, and index 2 primers. These are loaded into a NextSeq 550 High Output 75-cycle kit and sequenced on the machine with settings: read 1 (custom): 20 bp; read 2 (disabled); index 1 (custom): 8 bp; index 2 (custom): 8 bp, and a custom Illumina high output recipe that initiates the run with 2 dark cycles, before acquiring read data.

Sequencing run folders were demultiplexed with bcl2fastq2 with the settings --minimum-trimmed-read-length 19 and --mask-short-adapter-reads 19, due to the small read length. Raw FASTQ files were converted to sgRNA count tables using a Python script slightly modified from Cui et al. (*70*). Count tables were processed using a custom R script derived from Rousset et al. (*71*) that treats sgRNA counts like RNA seq counts. These counts are fit to a negative binomial distribution and fold changes from pre-and post-dCas9 induction are estimated using DESeq2. Significant deviations in gene essentiality, compared to control, were identified by plotting experimental vs. control fold-change values by gene. A linear regression model is fit to this plot, and if removing a gene from the plot significantly increases the fit of the regression model, then it is called as a significant change.

### DNA damage assays

Selected strains were grown at 37°C and 250 rpm in LB Miller media to prepare an overnight starting culture. The next day, overnight cultures were diluted 1:100 to a final OD_600_ = 0.05 in LB Miller media and grown with shaking (250 rpm) until mid-log phase (OD_600_ = 0.5). Cells were harvested, washed 3 times with PBS, and resuspended in PBS to an OD_600_ of 0.5. Cultures were serially diluted in PBS and spotted on LB agar plates containing different genotoxic agents at concentrations indicated in the figure. Each condition was tested in four replicates.

### High-throughput magnetic tweezers

The magnetic tweezers apparatus is implemented on a custom-built inverted microscope described previously (*72–75*). Briefly, collimated LED illumination (NA = 0.79, Thorlabs, Germany) passes through the gap of a neodymium magnet pair (W-05-G, SuperMagnete, Switzerland), whose height and rotation are controlled by linear motors M-126-PD1 and C-150, with C-863 controllers (Physik Instrumente, Germany). The field of view is imaged by a CFI Plan Apochromat Lambda D 60× oil-immersion objective (Nikon, Germany) mounted on a PIFOC piezo stage (P-726, E-753 controller; Physik Instrumente, Germany). Flow-cell temperature is regulated through an objective-based heating system (*76*). The temperature is regulated within approximately 0.1 °C by connecting the foil to a PID temperature controller (TC200 PID controller, Thorlabs). Images are acquired by a CMOS camera (Dalsa Falcon2, Stemmer Imaging, Germany; 4096 × 3072 pixels, 6 µm pixel size). Instrument control was implemented in a custom Python software and real-time 3D bead tracking was performed using an open-source algorithm (available at http://www.github.com/jcnossen/qtrk). The detailed parameters for the tracking are provided in **Supplementary Method 1**.

### Flow cell assembly

Flow cells were assembled as described (*77*). Two glass coverslips (#1, 24 × 60 mm, Menzel GmbH, Germany) are separated by a double-layer Parafilm channel (∼40 µL volume). The bottom coverslip is coated with 0.3% w/v nitrocellulose in amyl acetate.

### DNA construct

The transcription template was derived from pIA1437 (**Table S2**). It comprises a region upstream of a T7A1 promoter followed by a stalling sequence, A29, i.e., a short DNA sequence that embeds adenosine only at positions 2 and 29. Following downstream, there is a portion of the *rpoB* region, interrupted by a region that comprises the sequence upstream of *waaQ* (with the native *ops* site, ∼500 bp from T7A1) and the first ∼100 bp of the gene. A second *ops* site, with an identical sequence, was inserted by site-directed mutagenesis by PCR ∼1.1 kbp downstream T7A1. The construct was designed with a digoxigenin handle 3 kbp downstream of the promoter, to apply force in the opposite direction with respect to transcription (opposing force, OF). The construct was assembled by restriction–ligation (*78*); PCR conditions and gel-purification details are provided in **Supplementary Method 2**.

### Stalled elongation complex preparation

The WT or Δω RNAP holoenzyme was loaded at the promoter site and stalled in elongation, similarly to the protocols previously presented in the literature (*79, 80*). Briefly, the RNAP holoenzyme (1 nM) was incubated with the DNA construct (1:1 molar ratio) in PURE-like buffer (50 mM HEPES, pH 7.5, 100 mM KGlu, 13 mM MgAc_2_, 2 mM spermidine, 1 mM DTT, 2 mM NaN_3_, 1 mg/ml BSA) supplemented with ATP, CTP, GTP (50 µM each), and ApU dinucleotide (100 µM) at 37°C for 10 min. Heparin was added to a final concentration of 100 µg/ml for 15 min to sequester the free RNAP. Aliquots were snap-frozen in liquid nitrogen and stored at −80°C.

### Surface preparation and single-molecule transcription assay

Polystyrene reference beads (1.1 µm) were incubated in the nitrocellulose-coated flow cell, followed by anti-digoxigenin antibodies (50 µg/ml, 40 min), high-salt wash (TE750 + 2 mM NaN_3_), and BSA passivation (10 mg/ml, 40 min) followed by a second step of high-salt wash. PBS + 2 mM NaN_3_ was used to wash away reference beads and equilibrate the flow cell after high-salt washes.

The following steps were all performed in NaCl TX buffer (20 mM Tris-HCl, pH 7.5, 150 mM NaCl, 13 mM MgCl_2_, 1 mg/ml BSA, 1 mM DTT, 2 mM NaN_3_). The stalled complexes were diluted at 1:40 and incubated for 30 min. After washing away the excess, MyOne beads (1 µm) were incubated, and non-specific interactions were removed by applying 5 pN through the channel and washing again. Tethers showing extension of 0.2 – 0.3 µm upon force cycling (0.1/7 pN) were selected, if not showing multiple tethering to a single bead. Transcription was restarted with NTPs (1 mM each, ATP/GTP/CTP/UTP) at 37°C in PURE-like buffer under constant applied force (7 pN) for 1.5 h.

### Data processing

Raw reference-subtracted traces were processed with custom Python software. Traces where RNAP restarted >10 min after NTP addition were excluded to focus on a more homogeneous population. Tracking artefacts were removed manually. Extension was converted to base pairs using the worm-like chain model (considering 0.34 nm bp^−1^ at full extension). Gaps were filled by linear interpolation with added Gaussian noise (1 SD), and traces filtered with a Kaiser–Bessel low-pass filter at 1 Hz.

## Supplementary Materials

The PDF file includes:

Supplementary text

Figures S1 to S6

Tables S1 and S2

## Acknowledgments

We thank Natacha Ruiz and Georgiy Belogurov for fruitful discussions, Jason Peters for the generous gift of bicyclomycin, and Mark Finazzo for constructing p454 plasmid.

## Authors contribution

IA, CH, and DD supervised research and acquired funding. BH, BW, LB and MC performed the experiments and analyzed the data. BW and IA wrote the first article draft. All authors have edited the manuscript.

## Data, code, and materials availability

All data needed to evaluate the conclusions in the paper are present in the paper and/or the Supplementary Materials. The RNA-seq data were deposited to GEO (GSE328708) and the CRISPRi screen sequencing data ─ to SRA (PRJNA1457489). RpoS deletion RNA-seq was taken from NCBI (GSE87856) (*18*). The Scripts for the CRISPRi screen are deposited to GitHub (https://github.com/BingWangK/omega-and-TRC). A link to the single-molecule analysis data and scripts will be provided upon acceptance of the manuscript.

