## Supplementary materials for "The ω subunit stabilizes transcribing RNA polymerase to balance processivity and collision resolution"

Barbare Khitiri *et al.*

**This PDF file includes:**

Supplementary Text
Figs. S1 to S6
Tables S1 and S2
References (1 to 9)

### Supplementary Text

**Supplementary Method 1**: Bead tracking algorithm

Bead tracking is performed in real time on the CPU. For each bead, a squared region of interest (ROI) of 60 pixels is defined, and a lookup table (LUT) is constructed by recording the diffraction-pattern profile over 100 axial steps of 50 nm. Axial displacement is determined by the Quadrant Interpolation (QI) algorithm using 3 iterations. Mechanical drift is corrected by an autofocus routine that repositions the piezo when the reference-bead displacement exceeds a 5 nm threshold, with a 2 s refresh interval. Data points outside the LUT range are discarded (replaced by NaN).

**Supplementary Method 2**: PCR conditions for DNA construct preparation

Stem fragment: amplified from pIA1437 by Phusion (New England Biolabs (NEB)) using HF buffer. Cycling: 20 s initial denaturation at 98°C; 35 cycles of 10 s at 98°C, 30 s at 66°C, and extension at 72°C; 10 min final extension at 72°C.

Handle fragment: amplified from λ bacteriophage DNA by Taq (NEB) using Thermopol buffer in the presence of digoxigenin-conjugated UTP. Cycling: 20 s at 95°C; 35 cycles of 10 s at 95°C, 30 s at 52°C, and extension at 68°C; 10 min final extension at 68°C.

Both reactions used a Proflex PCR machine (ThermoFisher). Primers are listed in **Table S2**.

After BsaI-HFv2 (NEB) digestion and heat inactivation, fragments were purified by spin column (NEB) and ligated with T4 DNA ligase (NEB) in the presence of 12.5% (w/v) PEG-6000 (ThermoFisher). The ligation product was gel-purified from 1% agarose / 0.5×TAE stained with SYBR Safe (ThermoFisher). Concentrations were determined by a Denovix DS-11 spectrometer (DeNovix) and gel images acquired on an Amersham ImageQuant 800 imaging system (Cytiva).

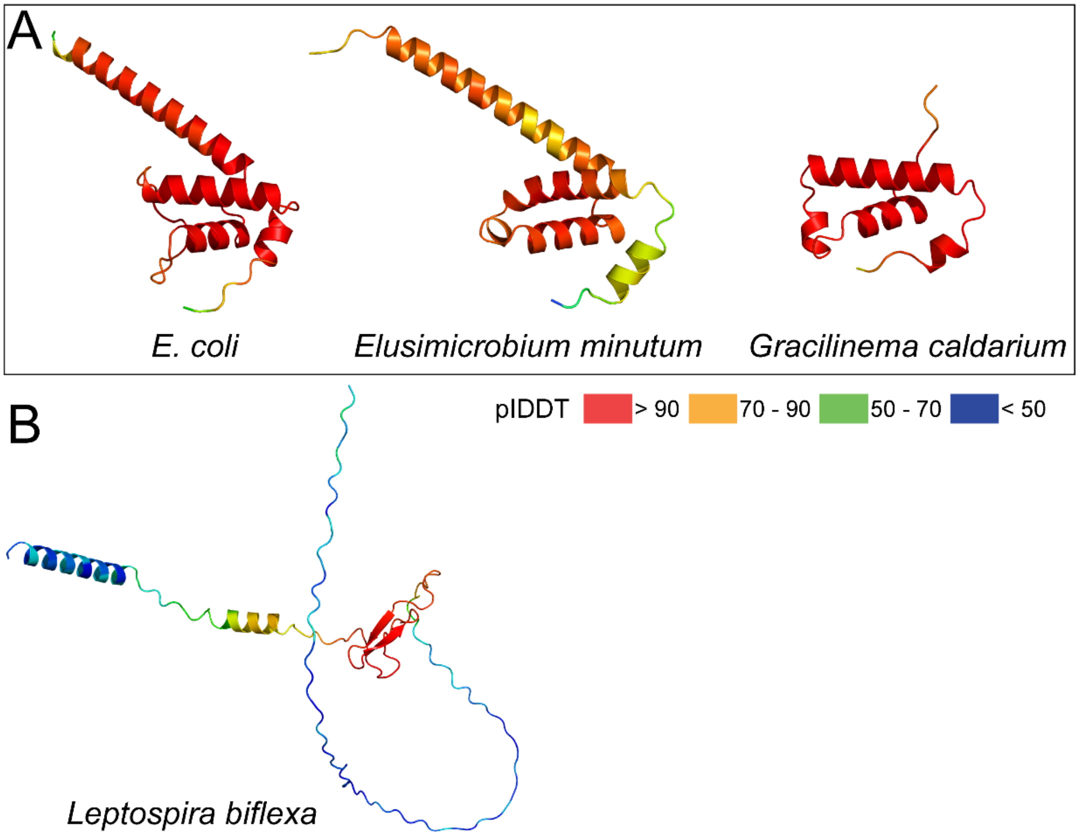


**Fig. S1. AlphaFold3 predicted structures of ω homologs.**
(**A**) The ω homologs (**Table S1**) from *E. minutum* (Elusimicrobiota) and *G. caldarium* (Spirochaetota), found by BLASTp, have similar structures to *E. coli* ω. (**B**) Predicted structure of an unknown protein (WP_012387502.1) from *Leptospira biflexa* (**Table S1**), which is located in the same locus as other Spirochaetota species (**Fig. 1C**).

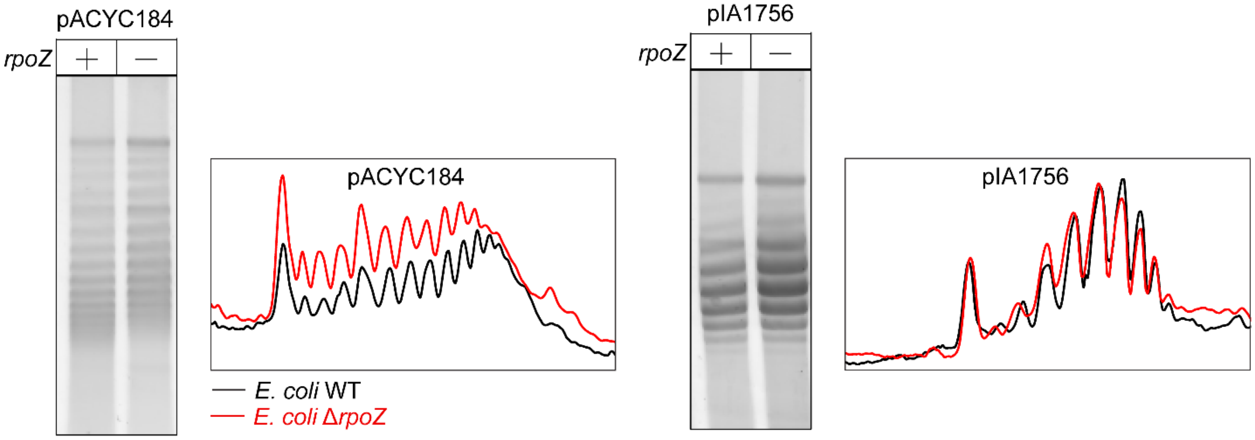


**Fig. S2. Effects of ω on plasmid topology.**
Two plasmids (**Table S2**) were tested in wild-type (*rpoZ*+) and Δ*rpoZ* (*rpoZ*-) strains. Plasmids were extracted from mid-log phase (OD = 0.4) *E. coli* cells. The distribution of topoisomers is plotted.

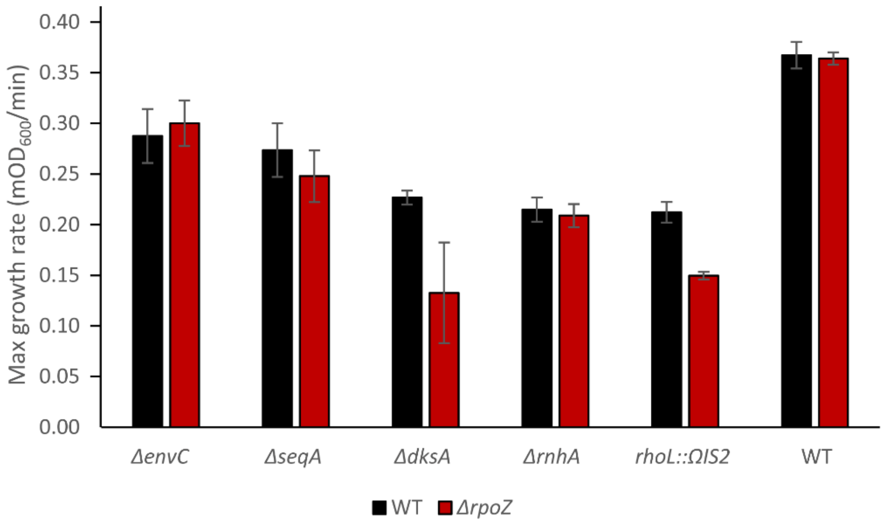


**Fig. S3. Growth assays of selected CRISPRi hits.**
Maximum growth rate ($mOD_{600}$/min) comparison between single mutants identified by CRISPRi screening in WT background (black bars) and double mutants in Δ*rpoZ* background (red bars). Error bars represent SD (Δ*envC,* n=3; Δ*rpoZ/rhoL::ΩIS2*, n=4; others, n=5).

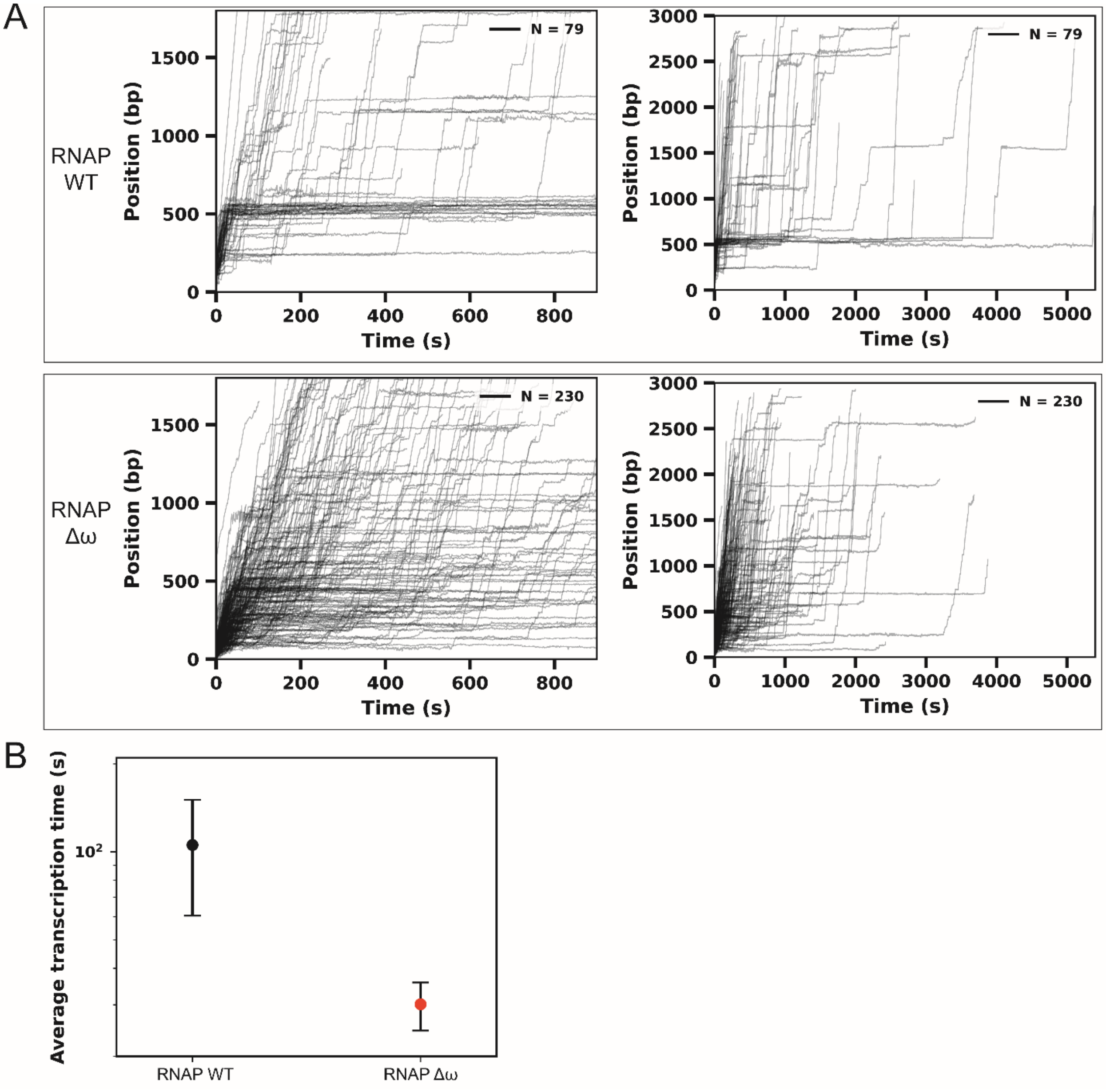


**Fig. S4. WT and Δω RNAP activity traces under 7 pN of applied force in OF configuration.** (**A**) Zoom-in of the first 15 min and 90 min of activity, in PURE-like buffer supplemented with 1 mM NTPs at 37°C, of the wild type (WT) and Δω RNAP. (**B**) Average transcription time for the WT and the Δω RNAP through the region of the *ops* site in the template (between 450 and 500 bp; **Fig. 4C**). The error bars display the standard deviation of the means obtained by 100 bootstraps procedure.

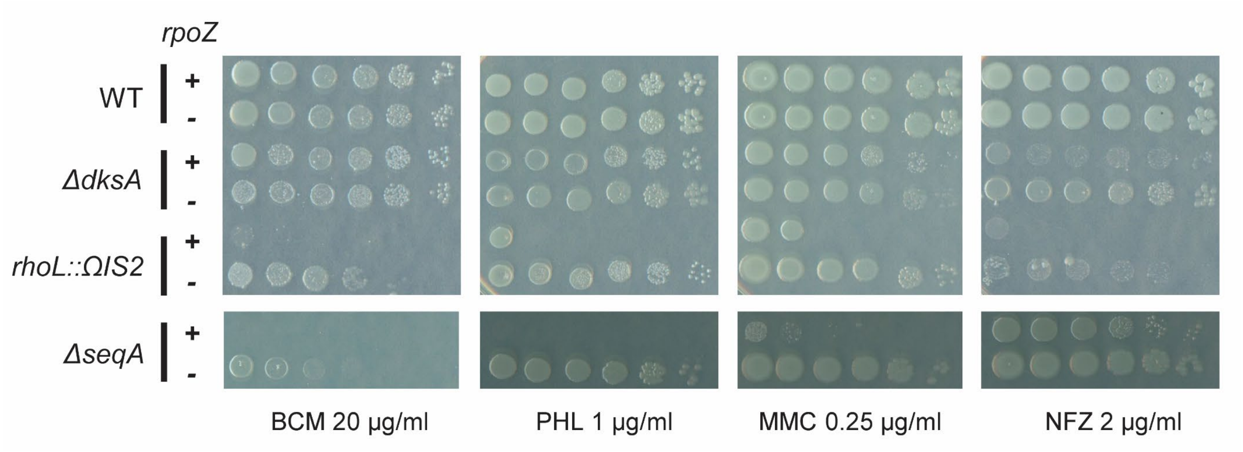


**Fig. S5. Effects of the *rpoZ* deletion on survival of Δ*dksA*, *rho*↓ and Δ*secA* under genotoxic stress.** Serial dilutions of cultures were plated on LB agar plates supplemented with the indicated DNA-damaging agents; see **Fig. 5**. NFZ, nitrofurazone.

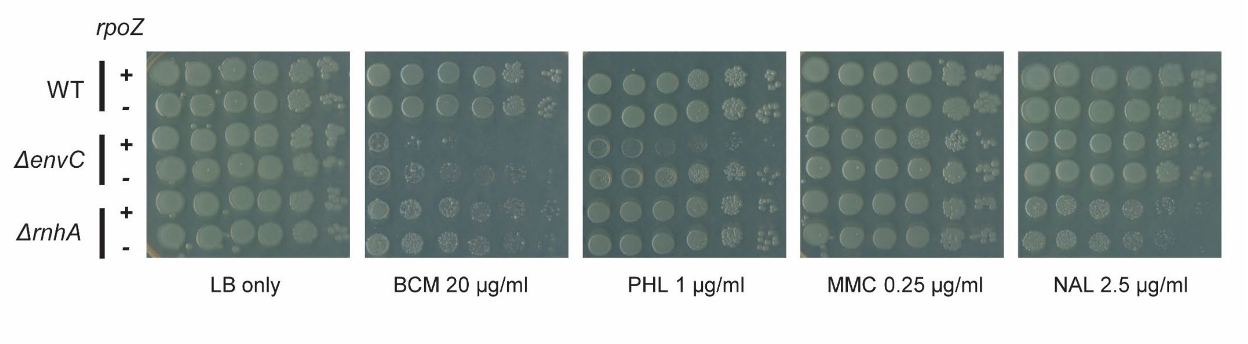


**Fig. S6. Effects of the *rpoZ* deletion on survival of Δ*envC* and Δ*rnhA* under genotoxic stress.** Serial dilutions of cultures were plated on LB agar plates supplemented with the indicated DNA-damaging agents.

**Table S1. The presence of ω homologs in four phyla was manually assessed using BLASTp**. For each phylum, a ω homolog from that phylum was used as the query sequence. Because there is no ω homolog identified in Elusimicrobiota by Annotree, a ω homolog from phylogenetically related Oederibacteriota (OGF51039.1) was used as a surrogate query. If a ω homolog was found by BLASTp, the NCBI protein ID will be shown. No, not found.

| Class | Order | Family | Genus | Species | Genome ID | Protein ID |
|---|---|---|---|---|---|---|
| Spirochaetota | | | | | | |
| Spirochaetia | Treponematales | Breznakiellaceae | Gracilinema | *Gracilinema caldarium* | GCF_000219725.1 | WP_013968654.1 |
| Spirochaetia | Borreliales | Borreliaceae | Borreliella | *Borreliella burgdorferi* | GCF_000008685.2 | WP_002557409.1 |
| Spirochaetia | Sphaerochaetales | Sphaerochaetaceae | Sphaerochaeta | *Sphaerochaeta globosa* | GCF_000190435.1 | WP_013607454.1 |
| Leptospiria | Leptospirales | Leptospiraceae | Leptospira_A | *Leptospira biflexa serovar Patoc* | GCF_000017685.1 | No |
| Leptospiria | Turneriellales | Turneriellaceae | Turneriella | *Turneriella parva* | GCF_000266885.1 | No |
| Brachyspiria | Brachyspirales | Brachyspiraceae | Brachyspira | *Brachyspira intermedia* | GCF_000223215.1 | WP_014487554.1 |
| Patescibacteriota | | | | | | |
| Patescibacteriia | Patescibacteriales | Patescibacteriaceae | Patescibacterium | Candidatus Falkowbacteria bacterium | GCA_016699775.1 | No |
| Saccharimonadia | Saccharimonadales | Saccharimonadaceae | Saccharimonas | Candidatus Saccharimonas aalborgensis | GCF_000392435.1 | No |
| Minisyncoccia | Minisyncoccales | Minisyncoccaceae | Minisyncoccus | *Minisyncoccus archaeiphilus* | GCF_047159735.1 | No |
| Gracilibacteria | Absconditabacterales | Absconditicoccaceae | Absconditicoccus | Candidatus Absconditicoccus praedator | GCF_021057185.1 | No |
| Chloroflexota | | | | | | |
| Anaerolineae | Anaerolineae | Anaerolineaceae | Anaerolinea | *Anaerolinea thermophila* | GCF_000199675.1 | No |
| Chloroflexia | Chloroflexales | Chloroflexaceae | Chloroflexus | *Chloroflexus aurantiacus* | GCF_000018865.1 | No |
| Dehalococcoidia | Dehalococcoidales | Dehalococcoidaceae | Dehalococcoides | *Dehalococcoides mccartyi* | GCF_000011905.1 | No |
| Ktedonobacteria | Ktedonobacterales | Ktedonobacteraceae | Ktedonosporobacter | *Ktedonosporobacter rubrisoli* | GCF_004208415.1 | No |
| Elusimicrobiota | | | | | | |
| Elusimicrobia | Elusimicrobiales | Elusimicrobiaceae | Elusimicrobium | *Elusimicrobium minutum* | GCF_000020145.1 | WP_012415008.1 |
| Endomicrobiia | Endomicrobiales | Endomicrobiaceae | Endomicrobium | *Endomicrobium proavitum* | GCF_001027545.1 | WP_052569726.1 |

**Table S2. Plasmids, strains, and oligonucleotides used in this study.**

| **Plasmids** | | |
|---|---|---|
| **Lab ID** | **Features** | **Source** |
| p216 | pACYC184: p15A origin; $Cm^R$ | R. Landick |
| P217 | T7A1-U less region (20 bp) – $\lambda_{TR2}$ terminator; $Amp^R$ | R. Landick |
| p316 | pCP20, $Amp^R$, $Cm^R$ | (*1*) |
| p453 | pKDsgRNA-NT; Spct [Addgene #89960] | Addgene |
| p454 | pKDsgRNA-p15; Spct | This study |
| pIA263 | $\lambda P_R$ promoter – C-less A26 ITR – T7Te terminator; $Amp^R$ | (*2*) |
| pIA264 | $\lambda P_R$ promoter – C-less A26 ITR – T3Te terminator; $Amp^R$ | This study |
| pIA265 | $\lambda P_R$ promoter – C-less A26 ITR – P14 terminator; $Amp^R$ | This study |
| pIA299 | T7 RNAP promoter – *lac* operator – *rpoA, rpoB, rpoC* - 6XHis – T7 terminator; $Amp^R$ | (*2*) |
| pIA692 | $\lambda P_R$ promoter – C-less A26 ITR – *rrnB* T1 terminator; $Amp^R$ | (*3*) |
| pIA447 | T7A1-U less region (37 bp); $Amp^R$ | This study |
| pIA1239 | $\lambda P_R$ promoter – C-less A26 ITR – NusA dependent *rsxC* terminator; $Amp^R$ | (*4*) |
| pIA447 | T7A1-U less region (37 bp); $Amp^R$ | This study |
| pIA1437 | T7A1-U less region (29 bp)-*ops* 1-*ops* 2; $Amp^R$ | This study |
| pIA1756 | pRSF1010 derivative; $Kn^R$ | (*5*) |
| pBK1 | pKDsgRNA-*rpoZ*; Spct | This study |

| **Strains** | | | |
|---|---|---|---|
| **Lab ID** | **Parent** | **Genotype** | **Source** |
| CH30 | - | *E. coli* MG1655 | CH Lab Stock |
| CH661 | - | *E. coli* BW25113 Δ*rpoZ*::Kan:FRT | Keio collection |
| CH7192 | LC-E75 | $attB_{186}$::$P_{tetA}$-dCas9::FRT, $att_{Lambda}$::mCherry::FRT | (*6*) |
| CH8186 | CH30 | $attB_{186}$::$P_{tetA}$-dCas9::Cam:FRT | (*7*) |
| CH8201 | CH8186 | $attB_{186}$::$P_{tetA}$-dCas9::FRT | (*7*) |
| CH8202 | CH8186 | $attB_{186}$::$P_{tetA}$-dCas9::FRT | (*7*) |
| CH13841 | CH8201 | $attB_{186}$::$P_{tetA}$-dCas9::FRT, ΔrpoZ::FRT | This study |
| CH13842 | CH8201 | $attB_{186}$::$P_{tetA}$-dCas9::FRT, ΔrpoZ::FRT | This study |
| CH13843 | CH8202 | $attB_{186}$::$P_{tetA}$-dCas9::FRT, ΔrpoZ::FRT | This study |
| CH13844 | CH8202 | $attB_{186}$::$P_{tetA}$-dCas9::FRT, ΔrpoZ::FRT | This study |
| IA161 | - | *E. coli* DH5α | NEB |
| IA549 | XJB λDE3 | *E. coli* B XJB λDE3 Δ*rpoZ*::FRT | (*8*) |
| IA765 | - | *E. coli* MG1655 | L. Freddolino |
| IA899 | IA765 | *E. coli* MG1655 *ΔlacI-ZYA*::Kan | This study |
| IA900 | IA765 | *ΔlacI-ZYA proC*$^+$ | This study |
| IA902 | IA900 | *ΔlacI-ZYA proC*$^+$*rpoZ*::Kan | This study |
| IA909 | IA902 | *ΔlacI-ZYA proC*$^+$*ΔrpoZ*::FRT | This study |
| IA911 | - | *E. coli* JW5646 *envC*::Kan | (*9*) |
| IA912 | - | *E. coli* JW0674 *seqA*::Kan | (*9*) |
| IA917 | - | *E. coli* JW0141 *dksA*::Kan | (*9*) |
| IA920 | - | *E. coli* JW0204 *rnhA*::Kan | (*9*) |
| IA923 | IA909 | *E. coli* MG1655 *proC*$^+$ *ΔlacI-ZYA ΔrpoZ envC*::Kan | This study |
| IA924 | IA909 | *E. coli* MG1655 *proC*$^+$ *ΔlacI-ZYA ΔrpoZ seqA*::Kan | This study |
| IA929 | IA909 | *E. coli* MG1655 *proC*$^+$ *ΔlacI-ZYA ΔrpoZ dksA*::Kan | This study |
| IA932 | IA909 | *E. coli* MG1655 *proC*$^+$ *ΔlacI-ZYA ΔrpoZ rnhA*::Kan | This study |
| IA934 | IA909 | *E. coli* MG1655 *proC*$^+$ *ΔlacI-ZYA ΔrpoZ rhoL*::ΩIS2 | This study |
| IA944 | IA900 | *E. coli* MG1655 *proC*$^+$ *ΔlacI-ZYA envC*::Kan | This study |
| IA945 | IA900 | *E. coli* MG1655 *proC*$^+$ *ΔlacI-ZYA seqA*::Kan | This study |

| | | | |
|---|---|---|---|
| IA946 | IA900 | *E. coli* MG1655 *proC*$^+$ *ΔlacI-ZYA dksA*::Kan | This study |
| IA947 | IA900 | *E. coli* MG1655 *proC*$^+$ *ΔlacI-ZYA rnhA*::Kan | This study |
| IA948 | IA900 | *E. coli* MG1655 *proC*$^+$ *ΔlacI-ZYA rhoL*::ΩIS2 | This study |
| IA1003 | IA765 | *E. coli* MG1655 *ΔrpoZ* | This study |
| IA1047 | - | *E. coli* JW5604 *envC*::Kan | (*9*) |

**Oligonucleotides**

| Lab ID | Sequence (5' – 3') | Purpose |
|---|---|---|
| **PCR primers for *in vitro* transcription templates** | | |
| fw OF handle | AAAACCTAAGAGACCGGAACCAAAGGATATTCAGACG | Single molecule assays |
| rv OF handle | AAAAGGATCCCGTGATGACCTCATTA | |
| fw OF stem | CAGTGAATTCGAGCTCGGTA | |
| rv OF stem | AATTGGTCTCTGCAAGGTCTTTCTTCGCCTGTTTG | |
| 44 | BIO-GGAGAGACAACTTAAAGAGA | The TEC stability assay |
| 1112 | GGAAGATGATCTTCCGGGGGCTT | |
| 17 | CGTTAAATCTATCACCGCAAGG | $\lambda P_R$ upstream |
| 256 | CAGTTCCCTACTCTCGCATG | $\lambda P_R$ and T7A1 downstream |
| 338 | GGAGAGACAACTTAAAGAGA | T7A1 upstream |
| 1665 | GGCCGCAGTATTGACTTAAAGTCTAACCTATAGGATACTTACAGCCAG | T7A1 DNA competitor |
| 1666 | GATCCTGGCTGTAAGTATCCTATAGGTTAGACTTTAAGTCAATACTGC | |
| **Primers for strain construction and validation** | | |
| 2421 | CGGATGTACTCAAAGCGGAC | PCR screen of *rho*::ΩIS2 |
| 2430 | TCGCCGAGAGTGATCAGCTC | |
| 2901 | CGCATGAGCCGCCAAAAG | *rpoZ*_F1 |
| 2904 | CTTTGTGATTAACGACGACCT | *rpoZ*_R90 |
| 3407 | gcacGCGTCGCGCTCGTCAGATGC | sgRNA for *rpoZ* |
| 3408 | aaacGCATCTGACGAGCGCGACGC | |
| 3409 | TCATGCCCAGTCATTTCTTCACCTGTGGAGCTTTTTAAGTTCACAAAGCGGG<br>TCGCCCTTGTATCTGTTTGAAAGCCTGA | sgRNA template |
| 3704 | AAAAATTGCTGGCGACGCTA | *rep*_R |
| 3705 | GTGCAGACCGTAATGTTCCAG | *dksA*_R |
| 3706 | ACGGATGTCCGCTTCATTTT | *envC*_R |
| 3707 | GGGTTATGGGACGGTGTAAT | *seqA*_R |
| 3708 | CGCATCGTTTATTGCGTGAAG | *rnhA*_R |
| 3709 | CGTTGGCTACCCGTGATATT | *neo*_F |
| **Primers for RT-qPCR** | | |
| 3566 | GCCCTGGGCGTAAAAGATCA | *waaQ* |
| 3567 | CAACCTGATAGCCTCGCTGT | |
| 3568 | AGCCAATGCAAGAAAAGGCG | *waaR* |
| 3569 | GTTAGTTTTGCGTCAGCCCA | |
| 3570 | ACCATCCCCACGATGAATGTT | *waaU* |
| 3571 | TGCAAGTCGTAGTTGGTGGAA | |
| 3490 | GGAGCATATGGCCTCGACTC | *ihfB* |
| 3491 | TCGCCAGTCTTCGGATTACG | |
| 3670 | CGTCATTAGCGACGATTGCG | *dps* |
| 3671 | TAGCTCTGGGGACCACTCAA | |
| 3672 | CAGACGAAGGTTTACCCGCA | *osmE* |
| 3673 | GCGGGTTGTACGGCTTATGA | |
| 3674 | CAGCCGCTCACGGTTATTTC | *katE* |

| | | | |
|---|---|---|---|
| 3675 | CCCTGAACGGTAGAGAAACG | | |
| 3676 | AGGTGATGGAAGAAGCACCG | | *rpoD* |
| 3677 | AGATTCCACGCTGGAAAGCA | | |
| 3325 | TCAACAGCATCTACATGATGGCC | | *rpoC* |
| 3326 | CGCCATCAGACCACGCATAC | | |
| 3599 | ACGCGAAGAACCTTACCTGG | | 16S rRNA |
| 3600 | TTCACAACACGAGCTGACGAC | | |
| **Primers for CRISPRi screen** | | | |
| **OC #** | **Purpose** | **Sequence (5' – 3')** | **Index** |
| OC1332 | PCR1 | GTGACTGGAGTTCAGACGTGTGCTCTTCCGATCT AAAGGACCCGTAAAGTGATAATGAT | - |
| OC1334 | PCR1 | TTCCCTACACGACGCTCTTCCGATCTATTACTCG GCACGCCCGTCGCTCAGTCCTAGGTATAATACTA | CGAGTAAT |
| OC1341 | PCR1 | TTCCCTACACGACGCTCTTCCGATCTTCCGGAGA GCACGCCCGTCGCTCAGTCCTAGGTATAATACTA | TCTCCGGA |
| OC1342 | PCR1 | TTCCCTACACGACGCTCTTCCGATCTCGCTCATT GCACGCCCGTCGCTCAGTCCTAGGTATAATACTA | AATGAGCG |
| OC1343 | PCR1 | TTCCCTACACGACGCTCTTCCGATCTGAGATTCC GCACGCCCGTCGCTCAGTCCTAGGTATAATACTA | GGAATCTC |
| OC1540 | PCR1 | TTCCCTACACGACGCTCTTCCGATCTATTCAGAA GCACGCCCGTCGCTCAGTCCTAGGTATAATACTA | TTCTGAAT |
| OC1541 | PCR1 | TTCCCTACACGACGCTCTTCCGATCTGAATTCGT GCACGCCCGTCGCTCAGTCCTAGGTATAATACTA | ACGAATTC |
| OC1542 | PCR1 | TTCCCTACACGACGCTCTTCCGATCTCTGAAGCT GCACGCCCGTCGCTCAGTCCTAGGTATAATACTA | AGCTTCAG |
| OC1543 | PCR1 | TTCCCTACACGACGCTCTTCCGATCTTAATGCGC GCACGCCCGTCGCTCAGTCCTAGGTATAATACTA | GCGCATTA |
| OC1333 | PCR2 | AATGATACGGCGACCACCGAGATCTACACTCTTT CCCTACACGACGCT | - |
| OC1335 | PCR2 | CAAGCAGAAGACGGCATACGAGATCGAGTAAT GTGACTGGAGTTCAGACG | ATTACTCG |
| OC1344 | PCR2 | CAAGCAGAAGACGGCATACGAGATTCTCCGGA GTGACTGGAGTTCAGACG | TCCGGAGA |
| OC1345 | PCR2 | CAAGCAGAAGACGGCATACGAGATAATGAGCG GTGACTGGAGTTCAGACG | CGCTCATT |
| OC1346 | PCR2 | CAAGCAGAAGACGGCATACGAGATGGAATCTC GTGACTGGAGTTCAGACG | GAGATTCC |
| OC1347 | PCR2 | CAAGCAGAAGACGGCATACGAGATTTCTGAAT GTGACTGGAGTTCAGACG | ATTCAGAA |
| OC1348 | PCR2 | CAAGCAGAAGACGGCATACGAGATACGAATTC GTGACTGGAGTTCAGACG | GAATTCGT |
| OC1349 | PCR2 | CAAGCAGAAGACGGCATACGAGATAGCTTCAG GTGACTGGAGTTCAGACG | CTGAAGCT |
| OC1350 | PCR2 | CAAGCAGAAGACGGCATACGAGATGCGCATTA GTGACTGGAGTTCAGACG | TAATGCGC |
| OC1544 | PCR2 | CAAGCAGAAGACGGCATACGAGATCTGATTAA GTGACTGGAGTTCAGACG | TTAATCAG |
| OC1545 | PCR2 | CAAGCAGAAGACGGCATACGAGATTAAGCAGT GTGACTGGAGTTCAGACG | ACTGCTTA |
| OC1546 | PCR2 | CAAGCAGAAGACGGCATACGAGATGAGCTACG GTGACTGGAGTTCAGACG | CGTAGCTC |
| OC1547 | PCR2 | CAAGCAGAAGACGGCATACGAGATAGAGAGGC GTGACTGGAGTTCAGACG | GCCTCTCT |
